# Chloramphenicol and gentamicin reduce resistance evolution to phage ΦX174 by suppressing a subset of *E. coli* C LPS mutants

**DOI:** 10.1101/2023.08.28.552763

**Authors:** Lavisha Parab, Jordan Romeyer Dherbey, Norma Rivera, Michael Schwarz, Jenna Gallie, Frederic Bertels

**Author notes:** Corresponding authors: **Lavisha Parab**. Address: Max Planck Institute for Evolutionary Biology, August-Thienemann-Straße 2, 24306 Plön, Germany;<u>; phone:</u>. **Frederic Bertels**. Address: Max Planck Institute for Evolutionary Biology, August-Thienemann-Straße 2, 24306 Plön, Germany.

## Abstract

Bacteriophages infect Gram-negative bacteria by attaching to molecules present on the bacterial outer membrane, often lipopolysaccharides (LPS). Modification of LPS can lead to resistance to phage infection. In addition, LPS modifications can impact antibiotic susceptibility, allowing for phage-antibiotic synergism. The evolutionary mechanism(s) behind such synergistic interactions remain largely unclear. Here, we show that the presence of antibiotics can affect the evolution of resistance to phage infection, using phage ΦX174 and *Escherichia coli* C. We use a collection of 34 *E. coli* C LPS strains, each of which is resistant to ΦX174, and has either a “rough” or “deep rough” LPS phenotype. Growth of the bacterial strains with the deep rough phenotype is inhibited at low concentrations of chloramphenicol (and, to a much lesser degree, gentamicin). Treating *E. coli* C wildtype with ΦX174 and chloramphenicol eliminates the emergence of mutants with the deep rough phenotype, and thereby slows the evolution of resistance to phage infection. At slightly lower chloramphenicol concentrations, phage resistance rates are similar to those observed at high concentrations; yet, we show that the diversity of possible mutants is much larger than at higher chloramphenicol concentrations. These data suggest that specific antibiotic concentrations can lead to synergistic phage-antibiotic interactions that disappear at higher antibiotic concentrations. Overall, we show that the change in survival of various ΦX174-resistant *E. coli* C mutants in the presence of antibiotics can explain the observed phage-antibiotic synergism.

*Classification*: Biological Sciences, Evolution

## Introduction

The rise of antibiotic-resistant bacteria is a global threat to public health (World Health Organization, 2014). To tackle this issue, several alternatives to antibiotics have been proposed (Czaplewski et al., 2016). Among the most promising of these are bacteriophages (phages), which have been successfully applied to a growing number of antibiotic-resistant bacterial infections (Schooley et al., 2017; Chan et al., 2018; Abedon, 2019b; Pirnay et al., 2022). However, as with antibiotics, a major reason for failure of phage treatments is the evolution of resistance. The success of phages as therapeutic agents depends on our ability to limit the emergence of phage-resistant bacteria.

### Reducing the evolution of phage-resistance in bacteria

One way to reduce the evolution of resistance is co-treatment of infections with specific combinations of phages and antibiotics (Coulter et al., 2014; Torres-Barceló et al., 2014; Chaudhry et al., 2017; Valério et al., 2017; Kebriaei et al., 2022). Under these conditions, bacteria must evolve resistance to both the phage and the antibiotic to evade elimination. Interestingly however, the efficacy of combined treatments is not always intuitive: examples of both synergistic and antagonistic interactions have been described. For example, phages NP1 and NP3 display synergistic effects with ceftazidime and ciprofloxacin when killing *Pseudomonas aeruginosa* biofilms (Chaudhry et al., 2017). Phages combined with antibiotics led to between a hundred-fold and thousand-fold higher killing rates, compared with phages alone. Contrastingly, phage SBW25Φ2 combined with streptomycin has been shown to increase rates of phage-resistance in *Pseudomonas fluorescens* populations (Cairns et al., 2017). While it is clear that the presence of antibiotics can affect the evolution of phage-resistance, thus far the underlying mechanisms remain largely unexplored. In this work, we provide a detailed example of how, and why, antibiotics can affect the course of phage-resistance evolution in bacterial populations.

### Mechanisms of phage-resistance in bacteria

A common mechanism by which bacteria become resistant to phage is through the modification of cell surface components, such as lipopolysaccharide (LPS) molecules (Hancock and Reeves, 1976; Mutalik et al., 2020; Romeyer Dherbey et al., 2023). LPS molecules are found on the outer membrane of all Gram-negative bacteria, and are the most common phage receptor of Gram-negative-infecting phages (Bertozzi Silva et al., 2016; Bio-Conversion Databank Foundation, 2021). Most *Escherichia coli* strains produce a “smooth” LPS phenotype, consisting of a lipid A domain, a core oligosaccharide (consisting of an inner-core and an outer-core), and a polysaccharide called the O-antigen (Frirdich and Whitfield, 2005). Some strains, such as *E. coli* C, lack the O-antigen and hence produce a “rough” LPS phenotype (Yethon et al., 1998). Further truncation of the LPS by removal of the outer-core oligosaccharide leads to a “deep rough” LPS phenotype (Nikaido and Vaara, 1985).

### E. coli C readily evolves resistance to ΦX174 through LPS alterations

Phage ΦX174 uses LPS to infect *E. coli* C (Michel et al., 2010). ΦX174 is one of the oldest and most widely-used phage model systems (Sinsheimer, 1959; Sanger et al., 1977; Smith et al., 2003). Surprisingly though, the mechanisms by which *E. coli* C can gain resistance to ΦX174 – and how these can be overcome – have only been explored recently (Romeyer Dherbey et al., 2023). In this previous study, we showed that *E. coli* C populations readily become resistant to ΦX174 infection through mutations that generate truncated LPS structures (**Figure 1**), and that ΦX174 can evolve to overcome this type of resistance.

**Figure 1.**
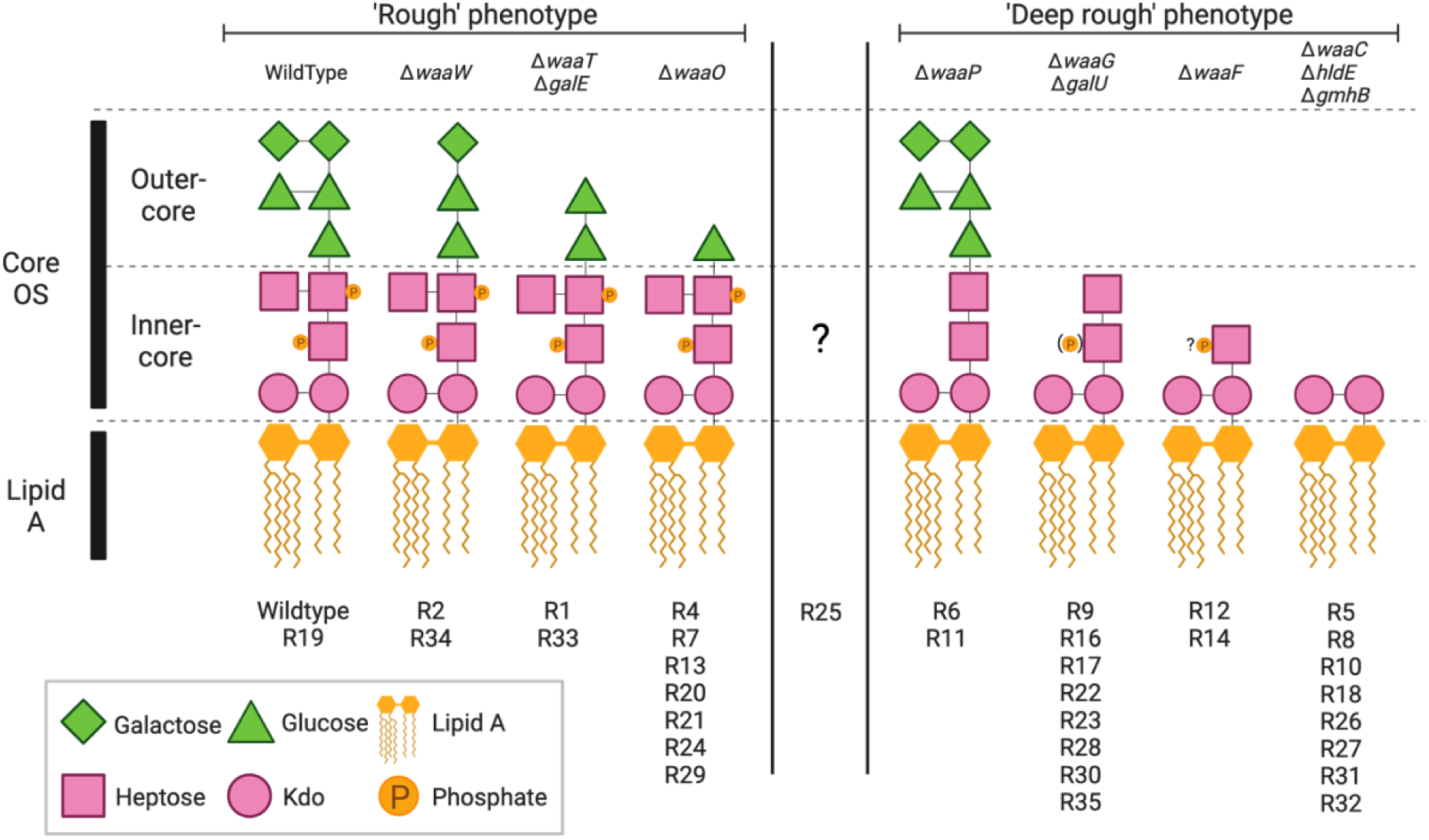
Predicted LPS phenotypes of 33 phage-resistant *E. coli* C strains. Strains (R#) are categorised into “rough” (longer; outer-core present but modified, inner-core unmodified) or “deep-rough” (shorter; outer-core absent and/or inner-core modified), based on mutation location. LPS phenotype cannot be predicted for R25 as the mutated gene, *rfaH*, controls the expression of the entire *waa* LPS operon (Klein and Raina, 2019). While the presence or absence of LPS outer core is commonly used to categorise LPS phenotypes, previous studies with gene deletion mutants show that absence of phosphate molecules in the inner-core also leads to a “deep rough” phenotype, as seen in R6 and R11 (Yethon et al., 1998). Kdo: 3-deoxy-D-*manno*-octulosonic acid; “?P”: No information was found on the phosphorylation of Hep(I) in the absence of *waaF*; (P): only 40 % of the hexose phosphorylation was observed (Yethon et al., 2000). OS: oligosaccharide. Figure adapted from Romeyer Dherbey et al., 2023.

### LPS phenotype affects sensitivity to both phage and antibiotics

In addition to affecting infection by phages, LPS phenotype can influence susceptibility to antibiotics. In *E. coli*, deep rough LPS mutants are more susceptible to membrane-targeting antibiotics such as colistin (Ebbensgaard et al., 2018; Burmeister et al., 2020). Resistance to many antibiotics that do not target the membrane can also be affected, because LPS modifications can also change membrane permeability (Nikaido and Vaara, 1985; Nikaido, 2003). For example, *E. coli* K-12 strains carrying a deep rough LPS structure are more susceptible to hydrophobic molecules such as novobiocin and spiramycin (Nikaido and Vaara, 1985; Nikaido, 2003). However, truncated LPS molecules can also lead to increased antibiotic-resistance. In *P. aeruginosa*, rough mutants (carrying mutations in *galU*) can increase antibiotic-resistance (El’Garch et al., 2007; Dötsch et al., 2009). Hence, only some combinations of LPS-targeting phages, antibiotic, and target pathogen, are expected to reduce the evolution of resistance.

In the current study, we investigate the effects of antibiotics on the evolution of ΦX174-resistance in *E. coli* C. We begin by measuring the susceptibility of a set of 33 ΦX174-resistant *E. coli* C strains (Romeyer Dherbey et al., 2023) to chloramphenicol and gentamicin. We find that *E. coli* C mutants with progressively shorter LPS molecules tend to be increasingly susceptible to both antibiotics (especially chloramphenicol). These results suggest that phage-antibiotic combination treatment prevents the growth of some antibiotic-resistant bacteria, and hence reduces the number and diversity of observable resistance mutations. The results of subsequent fluctuation experiments are consistent with this prediction; no *E. coli* C mutants with severely truncated LPS molecules were detected in a ΦX174+chloramphenicol combination environment. Finally, we use an *E. coli* C and ΦX174 co-evolution experiment in liquid media to show that, indeed, the ΦX174+chloramphenicol combination environment is the most efficient (of all combination treatments tested) at reducing phage-resistance evolution. Together, our results show that predicting phage-resistance rates in bacteria can be possible with surprisingly accuracy, using only very simple susceptibility assays.

## Results

### Phage-resistance mutations affect *E. coli* C susceptibility to chloramphenicol and gentamicin

The effect of LPS phenotype on antibiotic susceptibility is expected to depend on the antibiotic in question. Hence, in our first experiment, we investigated the effect of LPS phenotype on the susceptibility of *E. coli* C to two different antibiotics: chloramphenicol and gentamicin. These antibiotics inhibit protein synthesis via distinct mechanisms, respectively targeting the 50S and 30S ribosomal subunits (Lin et al., 2018). We quantified the growth of 34 *E. coli* C strains with various predicted LPS phenotypes (see **Figure 1**) in the presence of increasing – but nonetheless low – concentrations of chloramphenicol or gentamicin on agar plates. The results are presented in **Figure 2** and discussed in more detail below.

**Figure 2.**
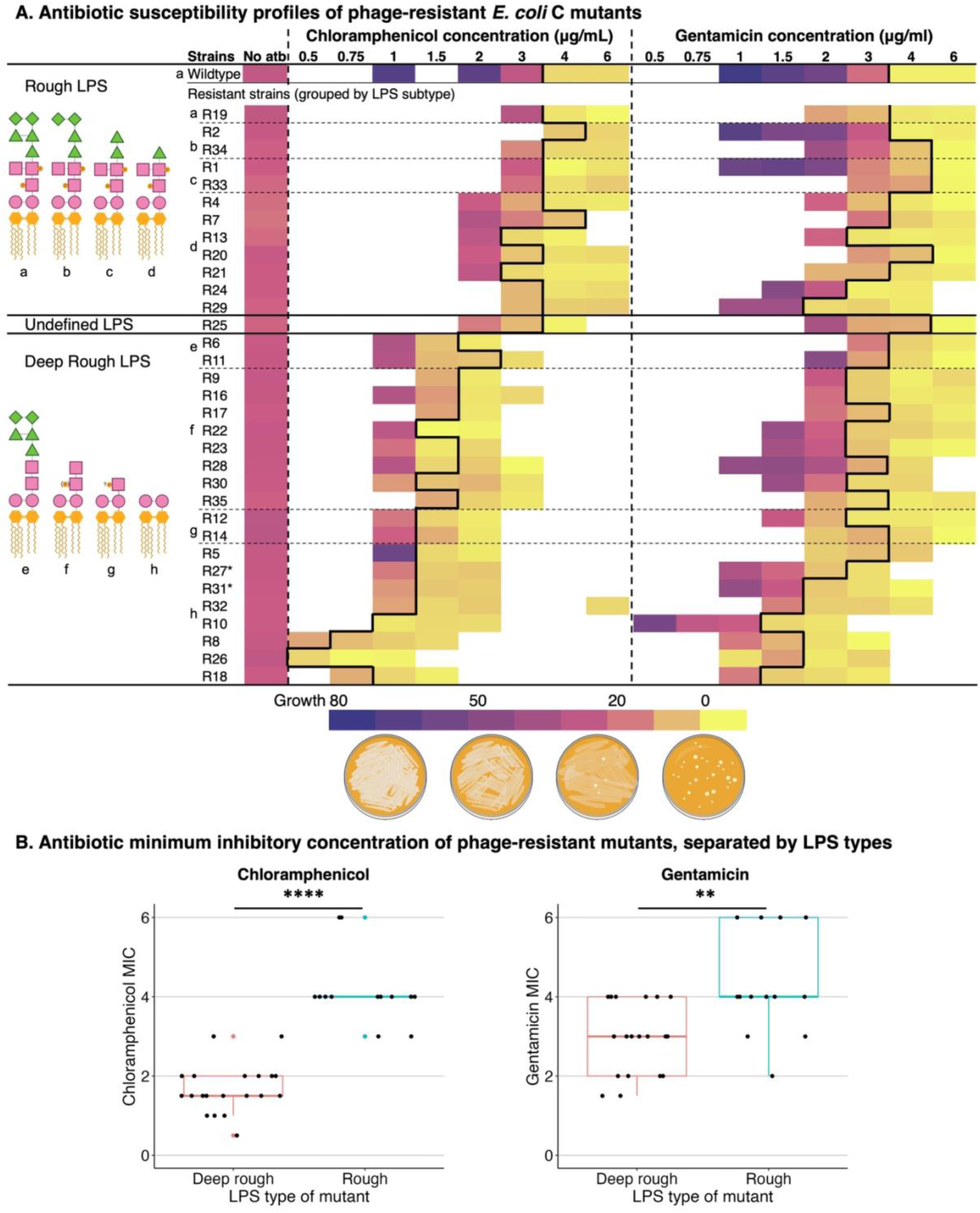
*E. coli* C deep rough LPS mutants show increased susceptibility to chloramphenicol and gentamicin. Heatmaps show growth of *E. coli* C wildtype and 33 *E. coli* C-derived, phage-resistant strains (R#) at increasing concentrations of chloramphenicol (left) and gentamicin (right). Growth is measured in 8-bit pixel greyscale values (see **Methods**) and decreases from violet (uniform lawn = greyscale values of 30 to 80) to yellow (no lawn = 0 to 8, and occasional colonies may be visible). White = not measured. Average greyscale values above 8 are defined as ‘growth’ (left of bold black border), while values below 8 are defined as ‘no growth’ (see **Methods**). Bacterial strains (rows) are ordered according to their predicted LPS phenotype (top to bottom: least to most truncated), and grouped as rough, undefined, or deep rough. Dotted horizontal lines group strains that have the same predicted LPS structure (shown on the left and indicated by letters a-h). *R27 and R31 have the same genotype (see **Supplementary Note 1**). The minimum inhibitory concentration (MIC) of *E. coli* C wildtype is ∼4 µg/mL for both antibiotics. Increased bacterial growth at sub-MIC antibiotic concentrations is observed in some cases; similar observations have been described previously (Frost et al., 2018; Mahrt et al., 2021; Farr et al., 2023; Aduru et al., 2024). Raw values plotted in **Supplementary Figures S1-S3**; raw data for this panel are available in **Supplementary Table S1**. (**B**) Boxplots show the MIC of the phage-resistant *E. coli* C strains in chloramphenicol (left) and gentamicin (right). The average MIC of strains predicted to carry rough (longer) and deep rough (shorter) LPS molecules differs for both antibiotics (*t*-test *p=*4.15E-08 and 0.002, respectively; Wilcoxon rank sum test for tied datasets *p=*1.05E-08 and 0.005, respectively). Bacterial strains predicted to display rough LPS are, on average, more susceptible to chloramphenicol and gentamicin.

### Chloramphenicol

The degree to which chloramphenicol affects *E. coli* C growth depends on LPS phenotype: progressively truncated LPS structures lead to increasing chloramphenicol susceptibility (Figure 2A**, Supplementary Figure S1**). More specifically, *E. coli* C strains displaying the rough LPS phenotype (including the wildtype) showed no detectable growth at chloramphenicol concentrations of >3-4 µg/mL, while the growth of strains displaying the deep rough LPS phenotype was already inhibited at chloramphenicol concentrations of 2 µg/mL or less (**Figure 2B, Supplementary Figure S3**). Further, a subset of deep rough mutants predicted to possess the shortest LPS structures (R8, R10, R18, R26) were most susceptible to chloramphenicol, with growth inhibition occurring at 1 µg/mL or lower.

### Gentamicin

Compared with chloramphenicol, susceptibility to gentamicin is less dependent on LPS phenotype. Many of the phage-resistant strains showed a similar growth profile to that of the wildtype across the gentamicin concentrations tested, with significant growth inhibition first occurring at concentrations of 3-4 µg/mL (Figure 2A-B**, Supplementary Figures S2 and S3**). One exception was a subset of six strains with severely truncated LPS phenotypes (*hldE*/*waaC* mutants: R8, R10, R18, R26, R31, R32), the growth of which was inhibited at lower levels of gentamicin (1.5-2 µg/mL) (see also **Supplementary Note S1**). Interestingly, the growth of five strains – namely, four strain with the rough LPS phenotype (R34, R1, R33, R20) plus the strain of unknown LPS phenotype (R25) – was first inhibited at a slightly higher gentamicin concentration than the wildtype (6 µg/mL versus 4 µg/mL). Hence, one might expect similar LPS mutations to be observed in gentamicin resistance screens. This is indeed the case for low-to-moderate levels of gentamicin (El’Garch et al., 2007; Ibacache-Quiroga et al., 2018).

The results in this section demonstrate that LPS phenotypes – and hence, phage-resistance genotypes – affect the susceptibility of *E. coli* C to both chloramphenicol and gentamicin. In general terms, shorter LPS structures tend to increase susceptibility to the antibiotic (particularly chloramphenicol).

### The presence of antibiotics is predicted to reduce diversity of phage-resistance mutations

In the previous section, we demonstrated that LPS mutations that decrease the susceptibility of *E. coli* C to ΦX174 can simultaneously increase susceptibility to low levels of antibiotics. This relationship suggests that the presence of antibiotics may affect the evolution of phage-resistance in *E. coli* C populations (and *vice versa*). Specifically, the frequency of phage-resistance mutations that render the bacterium more susceptible to antibiotic (i.e., those conferring the deep rough LPS phenotype), is expected to reduce in the presence of the antibiotic. In this section, we build a simple model to demonstrate this idea (model 1). This model can be outlined as follows:

1. Phage-resistance is primarily caused by deactivating mutation(s) in gene(s) encoding the LPS biosynthetic machinery (Romeyer Dherbey et al., 2023).
2. If we assume that these deactivating mutations occur uniformly between, and across the length of, each LPS gene, the total length of the gene(s) giving rise to phage resistance (i.e., the mutational target size) can be inferred.
3. Due to difference in which LPS phenotypes are permissible the mutational target size is affected by the presence – and concentration of – antibiotic. This information can be used to predict the relative rates of phage-resistance across environments.

*Inferring permissible mutational target sizes.* The mutational target size can be inferred from the genes that contain loss-of-function mutations in the 33 phage-resistant strains (Romeyer Dherbey et al., 2023). Mutations in a total of 14 genes, with a combined length of 11,736 bp, led to phage-resistant mutants (**Table 1**). This mutational target size is specific to the ΦX174-only environment; as the antibiotic concentration increases, the number of genes in which phage-resistance mutations are accessible (i.e., lead to survival) decreases (see **Figure 2**). Such ‘inaccessible’ genes can be deducted from the permissible target size in order to estimate the frequency of resistance to phage in each environment (**Table 1**).

**Table 1.**
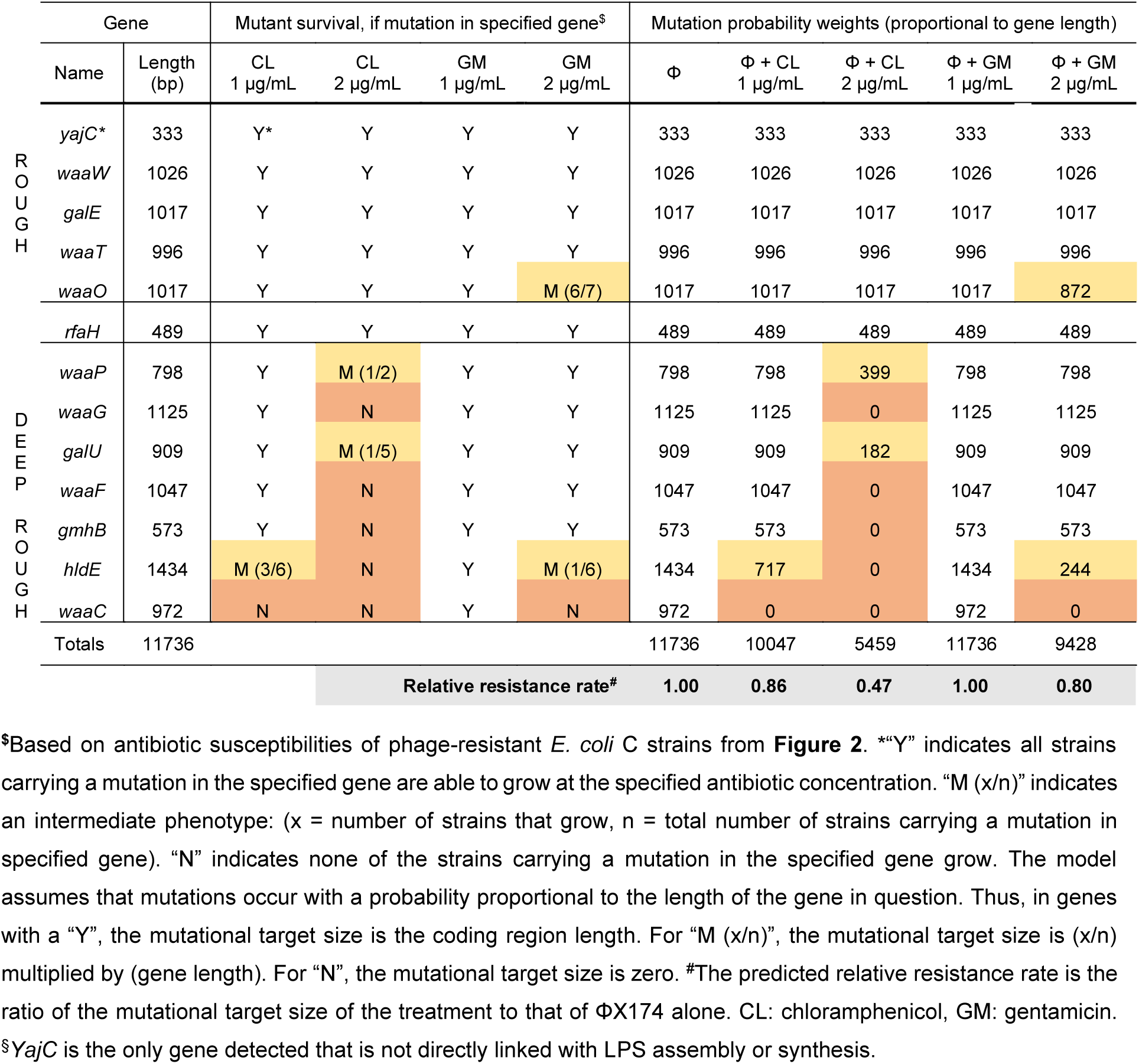
*A priori* prediction of relative phage-resistance rates in various environments (model 1).

The predicted relative resistance rates are provided in the final row of **Table 1** (and graphically in **Figure 3B**, dark red points). Overall, as antibiotic levels increase, the relative rate of phage-resistance is predicted to decrease. Most notably, the phage-resistance rate in 2 µg/mL chloramphenicol is predicted to be less than half (47 %) of that expected when treating the bacteria only with ΦX174. Combination treatment with gentamicin reduces the predicted resistance by only 20 % at the highest concentration tested (2 µg/mL).

**Figure 3.**
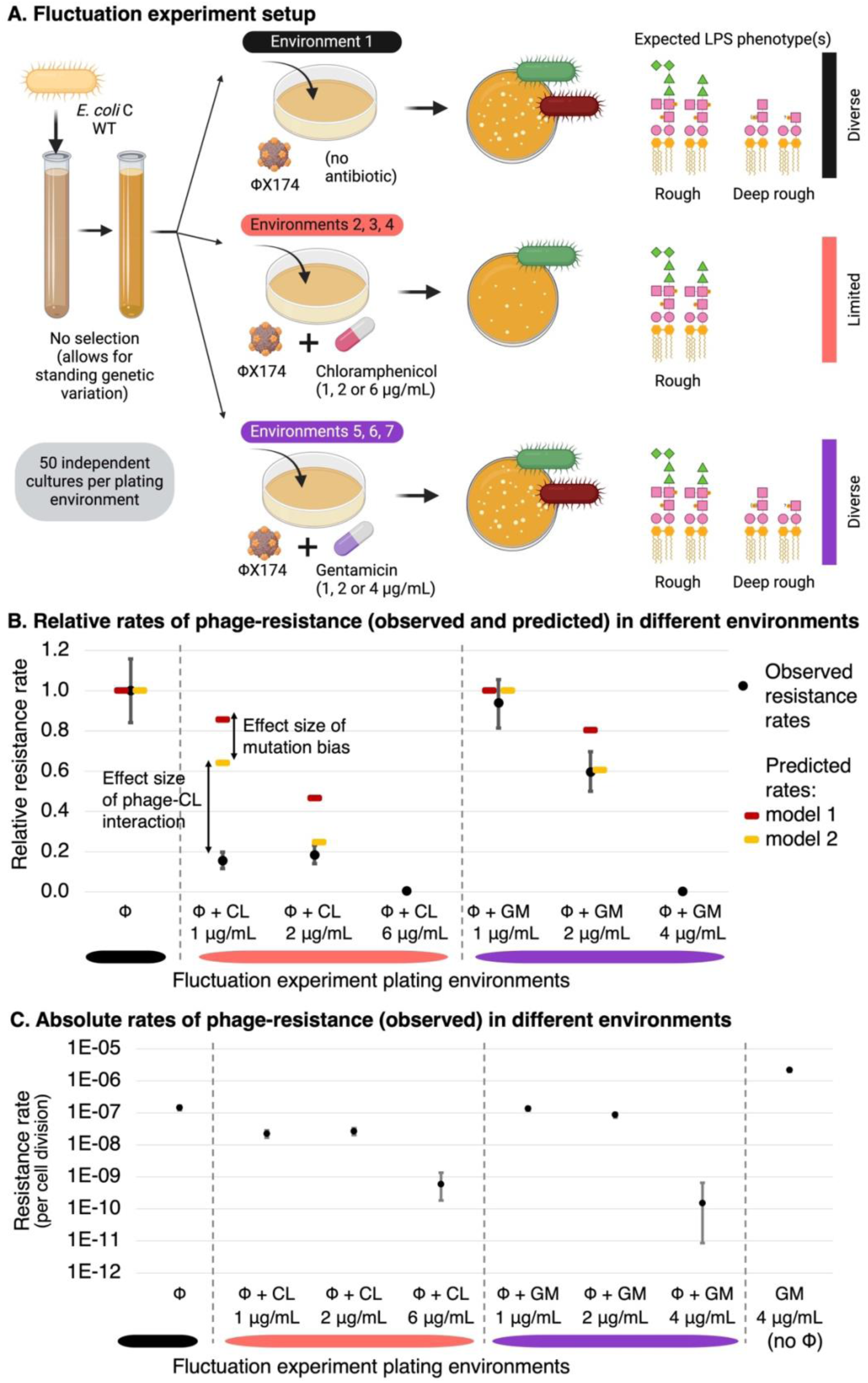
Antibiotics reduce the rate of resistance to ΦX174. **(A)** Schematic of the fluctuation test and expected results. Adding chloramphenicol at subinhibitory concentrations is predicted to reduce rates of resistance to ΦX174, due to inhibition of the growth of deep rough LPS phenotypes. Gentamicin, however, is predicted to have a much lower effect on the rate of resistance to ΦX174. (**B**) Resistance rates (and 95 % confidence intervals) for each phage+antibiotic plating environment, relative to the resistance rate in the ΦX174-only environment. Mean observed rates (black) are qualitatively in line with the predicted rates from model 1 (gene length model, dark red; derived in **Table 1**). Rates predicted by model 2 (gene proportion model inferred from the occurrence of mutations in a phage-only fluctuation test (see below), yellow; derived in **Supplementary Table S6**) are within 95 % confidence intervals for gentamicin (GM), and 2 % away from the 95 % confidence interval in 2 µg/mL chloramphenicol (CL). Resistance rates are not predicted at antibiotic concentrations inhibitory for wildtype (6 µg/mL CL, 4 µg/mL GM). The difference between models 1 and 2 can be mostly attributed to mutation bias (see text). The difference between model 2 and the observed resistance rate is most likely due to the effect size of phage-antibiotic interactions. (**C**) Absolute resistance rates (per cell division, and 95 % confidence intervals), including gentamicin at wildtype MIC without ΦX174. Rates of ΦX174 with antibiotic at inhibitory concentrations seem to be zero in panel B but are actually ∼400-1000-fold lower, as can be seen in panel C. In panel C, some 95 % confidence intervals are too small to be readily visible. Resistance rate values are provided in **Supplementary Table S2**, and raw colony counts are in **Supplementary Table S3**.

### Observed phage-resistance rates are qualitatively in line with *a priori* predictions

In order to test the relative phage-resistance rates predicted in the *a priori* model above, a classic fluctuation experiment was performed (Luria and Delbrück, 1943; Rosche and Foster, 2000). A total of 430 independent *E. coli* C wildtype populations were grown (without phage or antibiotics). Each grown population was spread onto LB agar plates containing either (set 1) ΦX174 alone, (set 2-4) ΦX174 in combination with chloramphenicol (at 1 µg/mL, 2 µg/mL, or 6 µg/mL), (set 5-7) ΦX174 in combination with gentamicin (at 2 µg/mL, 2 µg/mL, or 4 µg/mL), a total of 50 replicates for each of the seven environments (**Figure 3A**). In addition, 6 µg/mL chloramphenicol (set 8, n=30) and 4 µg/mL gentamicin (set 9, n=50) were used as control environments. For each plating environment, the number of resistant bacterial colonies arising from each population were counted. The distributions of colony numbers were used to calculate the observed relative phage-resistance rates (Zheng, 2017) (**Figure 3B-C**).

In model 1, the addition of chloramphenicol was predicted to reduce the rate of phage-resistance in *E. coli* C (**Table 1**). Indeed, the addition of chloramphenicol did significantly reduce relative phage-resistance rates (Likelihood ratio test (LRT) *p*<10^-5^ for both 1 µg/mL and 2 µg/mL) (**Supplementary Table S2**). However, the observed relative rates of phage-resistance were far lower than expected: 15 % at 1 µg/mL (∼four-fold lower than the predicted 86 %) and 18 % (∼two-fold lower than the predicted 47 %) (**Figure 3B**). The addition of gentamicin was predicted, and observed, to have no effect on phage resistance rates at 1 µg/mL gentamicin (LRT *p*=∼0.53). At 2 µg/mL gentamicin, the relative phage-resistance rate was reduced to 60 %, somewhat further than was predicted (80 %) (**Table 1**).

Overall, the addition of low concentrations of chloramphenicol or gentamicin did reduce the rate of phage-resistance in *E. coli* C (as predicted by model 1). However, the presence of antibiotics reduced the rate of phage-resistance much more effectively than predicted by the model. This could indicate that the reduction in rates of phage-resistance is not solely attributable to smaller mutational target sizes, and that other factors should be considered for more accurate predictions. However, before considering additional factors, it is important to rule out the possibility that minor improvements to the existing model may lead to a better match with the observations.

### Chloramphenicol skews mutations towards genes causing a rough LPS phenotype

So far, the results have demonstrated that combining ΦX174 with chloramphenicol (and, to a lesser degree, gentamicin) reduces the rate of phage-resistance in *E. coli* C. In model 1, we hypothesized that this is – at least partly – due to a reduction in the number of accessible mutations that lead to phage-resistance. To directly test this hypothesis, 222 phage-resistant *E. coli* C mutants were isolated from five of the fluctuation test plating environments, and each subjected to whole genome re-sequencing. A total of 239 mutations, belonging to a range of mutational classes, were identified (**Figure 4**, **Supplementary Figure S4**, **Supplementary Table S4**). Of the 222 mutant strains, 194 were found to contain mutations in the 14 previously identified LPS genes (Romeyer Dherbey et al., 2023), and two new LPS genes were identified as mutational targets (*hldD* and *gmhA*). Some degree of parallel evolution (identical mutations) was observed, both within and between plating environments (**Supplementary Note 2**). We note that, in four mutant strains, no putative mutations were identified (possibly due to low coverage of the LPS biosynthetic operon, see **Supplementary Table S4**).

**Figure 4:**
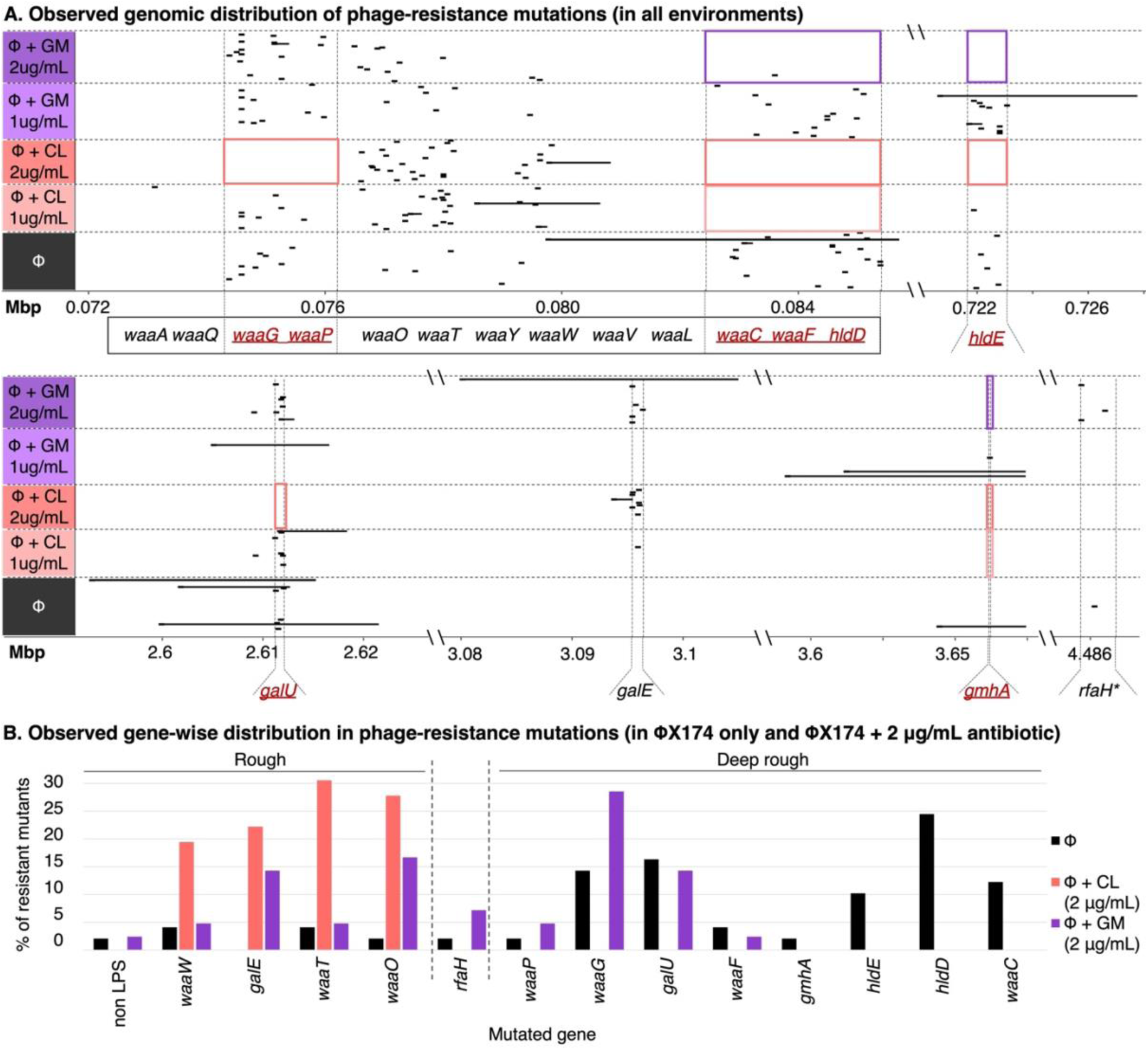
The presence of antibiotics reduces mutational target size for phage-resistance evolution in *E. coli*. **C.** (**A**) Genomic distribution of phage-resistance mutations in *E. coli* C, under five plating environments of the fluctuation test (rows). Large deletions (>100 bp) are indicated by solid horizontal lines scaled to the deleted region; all other classes of mutations are indicated as dashes. Red boxes indicate loci in which mutations are absent (or extremely rare) under some environments. Genes contributing to LPS inner core synthesis are underlined and in red; loss-of-function mutations in these genes are predicted to give a deep rough LPS phenotype. *RfaH is responsible for regulation of the LPS operon, and *rfaH* mutations can lead to a mixture of rough and deep rough types (Pradel and Schnaitman, 1991; Klein and Raina, 2019). (**B**) Gene-wise distribution of the phage-resistance mutations observed in the absence of, and presence of 2 µg/mL, antibiotics. Bars show the proportion of phage-resistant *E. coli* C mutants with a mutation in the specified gene (ΦX174-only n=49, ΦX174+CL(1 µg/mL) n=39, ΦX174+CL(2 µg/mL) n=36, ΦX174+GM(1 µg/mL) n=46, ΦX174+GM(2 µg/mL) n=42). CL: chloramphenicol, GM: gentamicin.

### Combining phage and antibiotics biases mutations towards rough LPS genes (Figure 5A)

Mutations in strains resistant to phage treatment alone are predominantly found in deep rough LPS genes (∼86 %), and only about ∼12 % are in rough LPS genes. As expected from our predictions, adding chloramphenicol enriches for rough mutants. At 1 µg/mL chloramphenicol the proportion of rough mutants increases from ∼12 % to ∼62 %, and the proportion of deep rough mutants decreases from ∼86 % to ∼36 %. At a concentration of 2 µg/mL chloramphenicol, all mutants are rough mutants. This result was somewhat surprising, since we predicted to observe at least some deep rough mutants (∼11 %) (**Figure 5A**). Gentamicin treatment also leads to a smaller proportion of deep rough mutants, but as expected from our predictions, the effect is much smaller. At a concentration of 1 µg/mL gentamicin deep rough mutants remain largely stable at ∼85 % and ∼15 % of the mutants are rough. At 2 µg/mL gentamicin, the ratio becomes more skewed towards a rough LPS with about 50 % deep rough mutants and ∼43 % rough mutants (**Figure 5A**).

**Figure 5.**
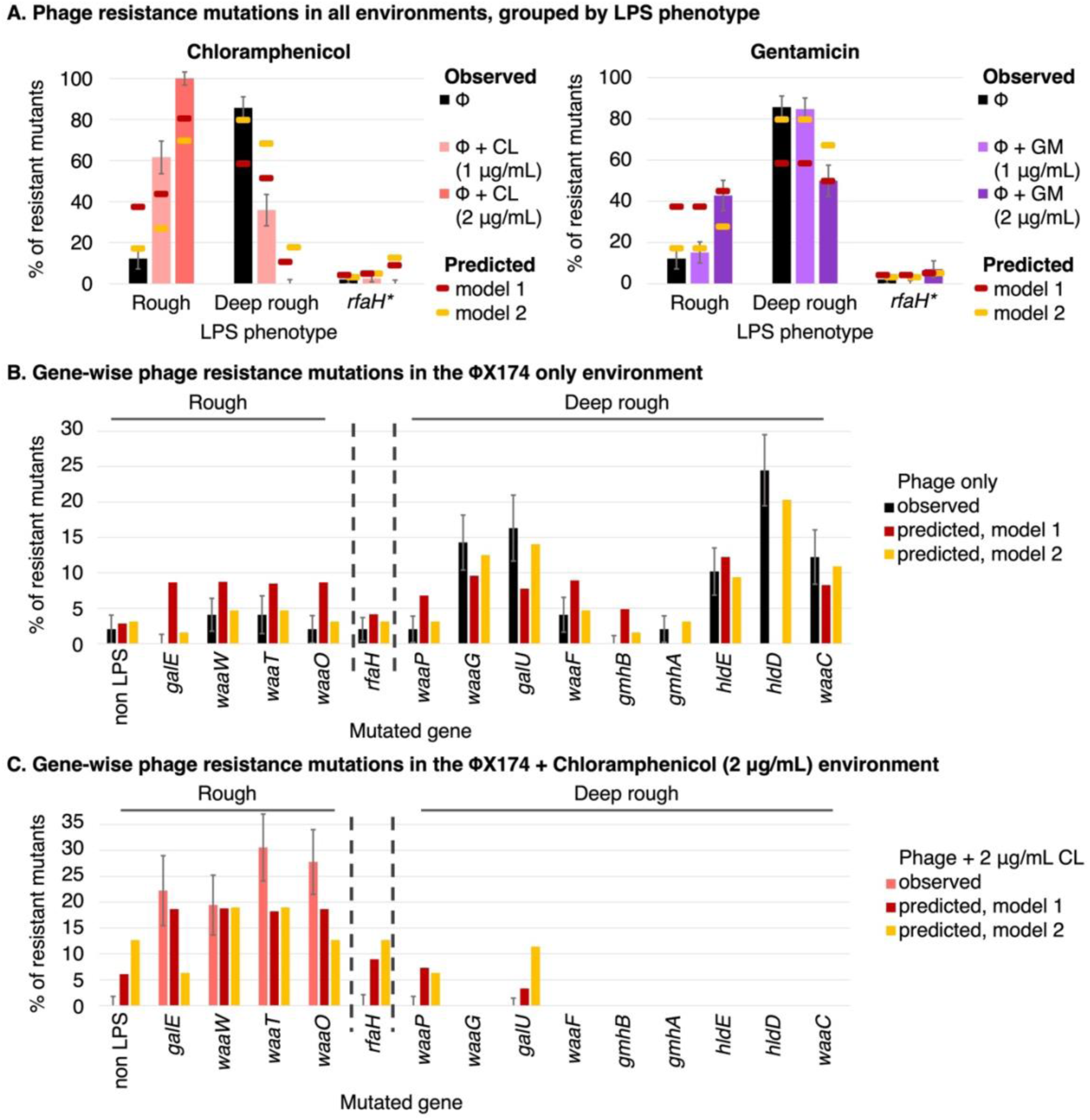
Chloramphenicol heavily skews phage-resistance mutations towards rough LPS loci. (**A**) Bars show the percentage of phage-resistant *E. coli* C mutants that carry mutations predicted to give rise to the rough or deep rough LPS phenotype, under various fluctuation test plating environments. Error bars show the standard deviation for each LPS type of 100 bootstrap replicates (resampling with replacement, see **Methods**). Model 1 predictions (dark red points) assume that the probability of mutation is proportional to gene length (**Table 1**). In model 2 (dark yellow points), the probability of mutation is based on the observed proportions in the ΦX174-only environment (**Supplementary Table S6**). Predictions for ΦX174+antibiotic environments are based on the susceptibility data (see **Figure 2**). Mutations in *rfaH* can affect any number of inner or outer core genes, and hence are in a separate LPS phenotype category. (**B**) Gene-wise distribution of phage*-*resistance mutations, in the absence of any antibiotics. The proportions predicted by model 2 differ from the observed proportions in the ΦX174-only environment because we add a pseudo-count of 1 to counts for each of the genes, to allow statistical handling of zero values (see **Methods**). “Non LPS” refers to genes whose potential involvement in LPS biosynthesis, assembly, or regulation remains uncharacterized (four genes across all experiments, see **Supplementary Note S3**). Error bars show the standard deviation 100 bootstrap replicates for each gene (**B**-**C**). (**C**) Gene-wise distribution of phage*-*resistance mutations, in the ΦX174+chloramphenicol (2 µg/mL) environment. As outlined in Model 2, *rfaH, galU* and *waaP* mutants are predicted to survive, but none were observed. (See **Supplementary Figure S5** for model 3).

### Adjusted model improves predictions and highlights potential phage-antibiotic interactions

Before attempting to predict outcomes of combination environments, our model should be able to accurately predict outcomes in the ΦX174-only environment. In the ΦX174-only environment, deep rough mutants were observed at significantly higher rates (86 %) than predicted from the mutational target size (model 1, 58 %) (**Figure 5A**, **Supplementary Table S5**). One possible explanation for this mismatch is that the distribution of mutations is not actually proportional to gene length, as model 1 assumes. For example, mutational bias could lead to unexpectedly high (or low) rates of particular mutations (Lind et al., 2019; Horton et al., 2021). To investigate this possibility, we used the proportion of mutants observed in each locus, in the ΦX174-only environment of the fluctuation test (a pseudo-count of 1 was added to each gene, to avoid false negatives, see **Methods**) (**Supplementary Table S6**). The adjusted predicted proportions are shown in yellow in **Figure 5B** (model 2). Adjusting the baseline predictions should, in theory, improve the predictions for the combination environments.

Using model 2 quantitatively improves predicted resistance rates (see **Figure 3B**), especially for environments with 2 µg/mL antibiotic (and phage). The new predicted resistance rate for 2 µg/mL gentamicin (61 %) is now within the 95 % confidence interval and close to the measured resistance rate (60%, with 95%CI 50-70%), while the predicted resistance rate in the 2 µg/mL chloramphenicol environment (i.e., 25 %) now lies almost within the 95 % confidence interval (18 %, with 95%CI of 14-23 %). However, the model 2-predicted distribution of rough and deep rough mutants fits the observed distribution less well than the model 1 predictions do (**Figure 5A**). The only environment where model 2 improved predictions of LPS mutant distribution above those in model 1, is ΦX174+1 µg/mL gentamicin (the environment closest to the ΦX174-only environment, based on our initial susceptibility assay) (**Figure 5A**). Model 2 also allows us to more accurately predict the effect of phage-antibiotic interactions on resistance rates by more accurately accounting for mutation bias (as shown in **Figure 3B**). The only environment where phage-antibiotic interactions appear to play a significant role is in the ΦX174+1 µg/mL chloramphenicol environment.

Model 2 qualitatively predicts the shift towards rough mutants, but without quantitative accuracy (**Figure 5A**). Since some *waaP* and *galU* mutants grow at 2 µg/mL chloramphenicol in our susceptibility assays, deep rough *waaP* and *galU* mutants are predicted to occur at a rate of 6 % and 11 %, leading to a non-zero deep rough prediction (**Figure 5A** and **5C, Supplementary Table S6**). *RfaH* mutants are predicted to occur at a rate of 13 %, because they also grow well in our susceptibility assay. Yet, *waaP, galU* and *rfaH* mutants were not observed at 2 µg/mL chloramphenicol (**Figure 5C**). Similarly, at 1 µg/mL chloramphenicol, mutations are predicted in *rfaH*, *waaF*, *gmhB*, and *gmhA* but none were observed (**Supplementary Figure S5C**). At 2 µg/mL gentamicin, mutations are predicted in *hldE*, *gmhA* and *gmhB* (deactivation of either is predicted to result in the deep rough LPS phenotype), but none were observed (**Supplementary Figure S5D**).

There are at least three possible reasons for the above discrepancies between predictions and observations. First, sampling only 40-50 mutants per environment could under-sample the proportion of mutated LPS genes. Sampling more mutants in the ΦX174-only environment may improve the accuracy of predicted proportions (and prevent the need for the addition of pseudo-counts in our models). Second, biological interactions between antibiotics and phages (Comeau et al., 2007; Gu Liu et al., 2020) may lead to the over-representation of different LPS genes compared to the ΦX174-only environment. Third, it is possible that what is considered growth in our susceptibility assay is not sufficient to be detected in the fluctuation experiment.

We are planning to test hypotheses one and two in future studies, hypothesis three can be tested by adjusting the threshold for growth in our susceptibility assay. When we increase the threshold for growth from a greyscale value of 8 to 12 (model 3), phage-resistance rate predictions remain largely unchanged from those in model 2, with the exception of the 2 µg/mL gentamicin environment, where the prediction worsens (**Supplementary Figure S5A**). Predicted distributions across LPS genes do not change significantly for most environments (**Supplementary Figure S5B**, **Supplementary Tables S6-S7**), but model 3 does indeed improve the gene-wise predictions for phage+2 µg/mL antibiotic (**Supplementary Figure S5D-E**). With the higher threshold, *gmhB*, *gmhA* and *hldE mutants* are no longer predicted to grow at 2 µg/mL gentamicin (**Supplementary Figure S5D**), neither are *waaP* and *galU* mutants at 2 µg/mL of chloramphenicol (**Supplementary Figure S5E**), which matches our experimental observations. However, none of the model adjustments explain the discrepancy between the predicted and observed phage-resistance rates, or proportions of deep rough and rough mutants at 1 µg/mL chloramphenicol (**Supplementary Figure S5A-B**). Hence, this discrepancy is most likely the result of direct or indirect interactions between phages and chloramphenicol.

Interactions between phages and chloramphenicol are also indicated by statistical comparisons between observations and expectations according to different models (**Table 2**). If there is no statistical difference between model and observation, we assume that the model performs well. As the difference between model and observation becomes more significant, the model fits the observations less well. Of the three models, models 2 and 3 provide the most accurate predictions of rough and deep rough mutant proportions (**Table 2**), and phage-resistance rates (**Figure 3B, Supplementary Figure S5**). The opposite is true for gene-wise distribution predictions, since model 1 is closer to the observations than later models (with the exception of the ΦX174+1 µg/mL gentamicin environment). This indicates that gene length is a reasonable proxy for the likelihood of observing a mutation in a particular gene (bottom of **Table 2**). Hence, adjusting the mutation proportions to the ΦX174-only environment leads to fewer errors for resistance rate and LPS type predictions, but greater errors for gene-wise predictions.

**Table 2.** Chi-squared tests to check goodness of fit between observed and predicted mutant proportions

| **LPS type**<br>**(expected and observed proportions of deep rough and rough mutants, Figure 5A)** |  |  |  |  |
| --- | --- | --- | --- | --- |
| Treatment | Model 1 (added genes) | Model 2 | Model 3 | Model 1 |
| Phage | 0.0046 | 0.576 | 0.576 | 0.0005 |
| Phage + 1 μg/mL CL | 0.0368 | 6.35E-06 | 6.35E-06 | 0.0779 |
| Phage + 2 μg/mL CL | 0.0124 | 0.000388 | 0.01052 | 0.0124 |
| Phage + 1 μg/mL GM | 0.0084 | 0.429 | 0.429 | 0.0012 |
| Phage + 2 μg/mL GM | 0.7795 | 0.059 | 0.555 | 0.838 |
| **Gene-wise distribution**<br>**(expected and observed proportions of mutations in each gene)** |  |  |  |  |
| Treatment | Model 1 (added genes)<sup>^</sup> | Model 2 | Model 3 |  |
| Phage | 6.20E-05* | 0.9981 | 0.9981 |  |
| Phage + 1 μg/mL CL | 1.69E-05 | 7.15E-07 | 7.15E-07 | |
| Phage + 2 μg/mL CL | 0.0553 | 3.16E-08 | 3.51E-06 | |
| Phage + 1 μg/mL GM | 0.0091 | 0.01022 | 0.0102 | |
| Phage + 2 μg/mL GM | 0.0051 | 7.60E-05 | 0.0005 | |

\*Values shown are *p*-values obtained by comparing the observed distribution of mutants (gene-wise or LPS type) with the distribution predicted by the given model. <sup>^</sup>Before comparing model 1, two missing genes *hldD* and *gmhA*, were added to resistance genes based on the underlying gene length assumption, to avoid zero expectation for any category.

### Chloramphenicol addition leads to strongest synergism in co-evolving *E. coli* C-ΦX174 populations

To test whether our above predictions also hold in more complex environments, we set up a serial transfer co-evolution experiment. The experiment began with ten distinct *E. coli* C-ΦX174-antibiotic combinations (see **Figure 6A**). The growth of twelve replicate cultures in each environment was recorded for 24 hours (day 1), after which 2 % of each culture was transferred to fresh media and growth continued for a further 24 hours (day 2). The most influential factor in the growth of day 1 cultures was the presence of phages; as expected, cultures with phages grew significantly more slowly than those without phages (**Figure 6B-C**, left panels). However, even cultures containing phages showed considerable bacterial growth by 24 hours, suggesting the rapid emergence of phage-resistant bacteria (including the emergence of phage resistant genotypes). Indeed, on day 2, the presence of phages is no longer the most influential factor on growth of the bacterial population (**Figure 6B-C**, right panels). Instead, the growth profiles of day 2 populations group not by phage presence, but instead by antibiotic: cultures with no antibiotic grew most quickly, then those with gentamicin, and finally those with chloramphenicol. Antibiotics becoming the dominating factor in our evolution experiment at day 2 is strong evidence of rapid phage resistance evolution. Further, the course of evolution is affected by the presence of antibiotics: phage resistance emerges rapidly when no antibiotic is present, but considerably more slowly when gentamicin or chloramphenicol is present. On day 2 for high antibiotic concentrations there is no significant difference between the antibiotic only and the antibiotic-phage treatments except for the 1 µg/mL chloramphenicol environment.

**Figure 6.**
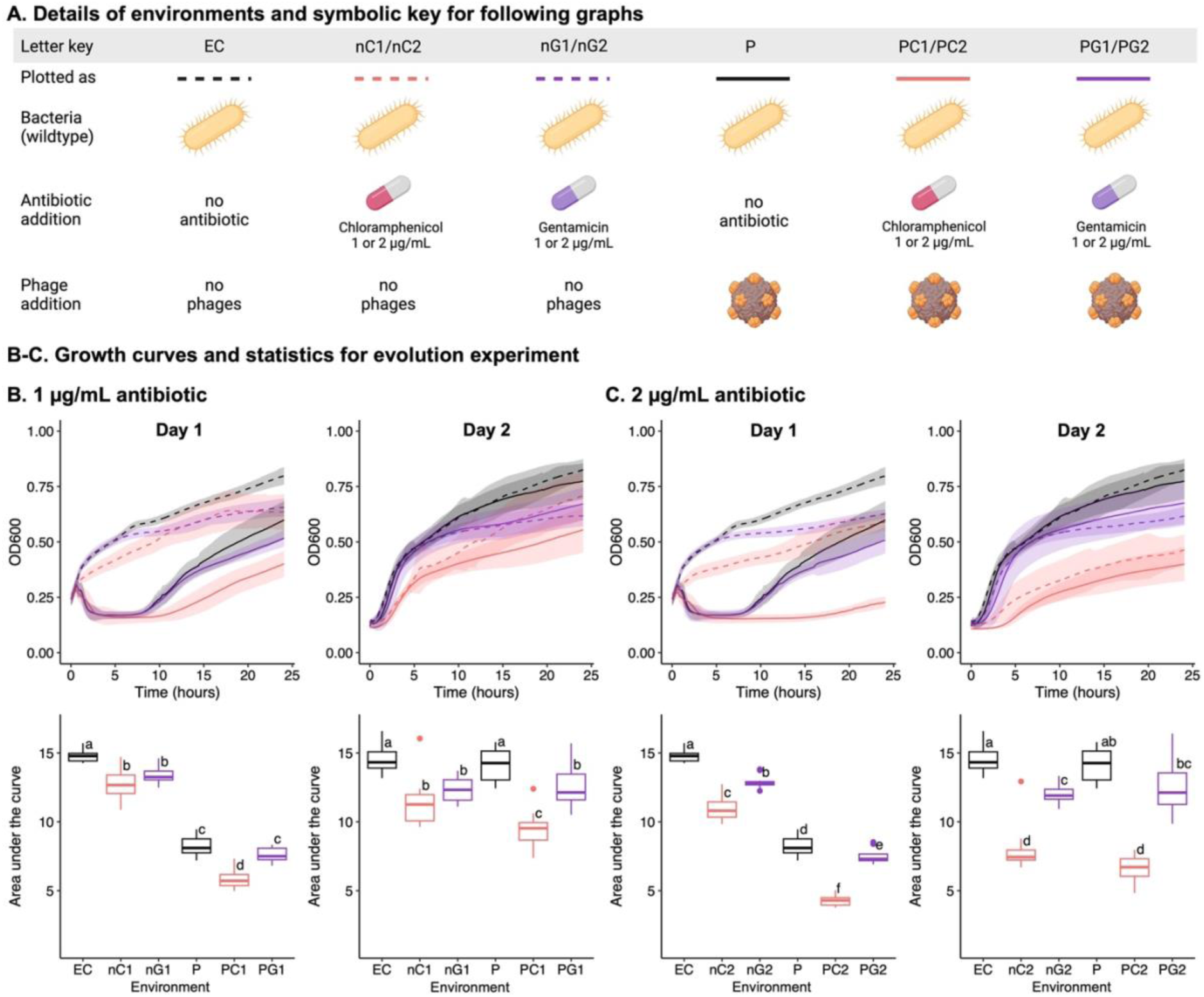
Chloramphenicol affects co-evolution of *E. coli* C and ΦX174. (**A**) Details of the 10 distinct *E. coli* C culture environments in the two-day evolution experiment, containing the indicated combinations of ΦX174, chloramphenicol, and/or gentamicin (n=12 per environment). Day 2 populations were founded by transferring 2 % of each population into fresh media under the same conditions as day 1. (**B-C**) The effect of 1 µg/mL (**B**) or 2 µg/mL (**C**) antibiotic on the growth of *E. coli* C cultures with and without ΦX174. Growth is presented as absorbance measures over time (top) and area-under-the-curve (AUC, bottom), on day 1 (left), and day 2 (right). Letters on the AUC plots denote statistical significance groups for each day (a-f): there is no statistically significant difference between environments that share a particular letter on one day, while there is a significant difference between environments that do not share the letter. For example, on day 1, in 1 µg/mL antibiotic (**B**), there is no significant difference between nC1 and nG1, or between P and PG1, but all other pair-wise combinations are significantly different (Tukey’s Honest Significant Difference test, see **Methods**). Statistical testing was done separately for (**B**) and (**C**).

Only for the 1 µg/mL chloramphenicol environment remain phages a significant growth inhibitor on day 2 (**Figure 6B**). This result is in line with the low level of phage resistance evolution at 1 µg/mL chloramphenicol in the solid fluctuation experiment (**Figure 3B**). Similarly, we posit that interactions between phage and antibiotic are likely the cause for the continued efficacy of phage treatment and prevent the emergence of fast growing phage resistant mutants.

## Discussion

Here we find that phage-antibiotic combinations can reduce the evolution of phage-resistance in bacteria, by inhibiting the growth of certain bacterial mutants. We start by quantifying the effect of chloramphenicol and gentamicin on the growth of 34 previously isolated ΦX174-resistant *E. coli* C strains. From these measurements, we predict that certain classes of mutations will not emerge in specified phage-antibiotic combination treatments, and hence these combination treatments will reduce the rate of phage-resistance evolution. We test these predictions by performing an experiment where we expose *E. coli* C wildtype to phage ΦX174 and antibiotics. The observed phage resistance rates generally agree with our predictions, and whole-genome sequencing of phage-resistant mutants support our predictions of mutational landscape accessibility. Overall, we show that antibiotics can reduce phage resistance evolution by preventing the growth of deep rough LPS resistance types. This effect is often referred to as an evolutionary trap, and can be useful for evolutionary steering (Baym et al., 2016; Barbosa et al., 2017; Burmeister et al., 2020). Interestingly, first order effects – effects which do not take interactions between phage and antibiotic into account – are largely sufficient to explain reduced phage resistance evolution in combination treatments. Finally, we show that our predictions hold even in a more complex co-evolution experiment, where phage-chloramphenicol combinations are the most potent treatments for slowing bacterial growth.

### Phage-chloramphenicol interactions likely amplify the effect of combination treatments

The original model assumption for the fluctuation assay posits that the outcome of the assay only depends on two parameters: the mutation rate and the mutational target size (Luria and Delbrück, 1943). The mutation rate should be identical across all treatments, since mutations occur during exponential growth in liquid culture in the absence of antibiotic and phages. The second parameter, the measured mutational target size, is very close to our expectations for 1 µg/mL chloramphenicol, and similar to the measured target size for 2 µg/mL gentamicin (**Figure 4A**). Yet, for gentamicin the predicted phage resistance rate (61 %) fits well with the observed resistance rate (60 %), while for chloramphenicol the rates differ significantly (64 % predicted versus 15 % observed, **Figure 3B**). Given that target size and mutation rates are identical between the environments, there must be other factors that explain our observations.

One possible explanation is that mutations do not guarantee resistance. It is possible that for 1 µg/mL chloramphenicol the mutations only provide a certain survival chance rather than guaranteed survival. Previous studies have shown that while low concentrations of chloramphenicol do not lead to stochastic population extinctions, combination of chloramphenicol and a bactericidal antibiotic does (Coates et al., 2018). Given that phages can be considered bactericidal agents, stochastic population extinctions could play a role in our experiment. If the stochastic extinction chance is around 50 %, then the expected resistance rate would correspond to our observed resistance rate. This stochastic survival effect may result from several possible factors, including physiological interactions between phage and chloramphenicol on the plate, local fluctuations in chloramphenicol concentration, or differences in physiological cell state (Akiyama and Kim, 2021).

The combination between a bactericidal agent (phage) and a bacteriostatic antibiotic (chloramphenicol) may lead to a trade-off explaining why phage resistance rates at 1 µg/mL and 2 µg/mL chloramphenicol are similar. At high chloramphenicol concentrations, the antibiotic can efficiently prevent the emergence of deep rough mutants. However, the efficiency of killing by the phage may be simultaneously compromised because of reduced bacterial growth. Slower growing bacteria inhibit phage infection and hence phage killing efficacy (Hadas et al., 1997; Coates et al., 2018). At slightly lower concentrations, chloramphenicol has a smaller effect on bacterial growth, rendering phage infection more efficient. These effects could conceivably balance each other out, leading to similar phage resistance rates at chloramphenicol concentrations of 1 µg/mL and 2 µg/mL.

Similar to our results in solid media, we also observe the strongest effect of combining phage and antibiotic in the 1 µg/mL chloramphenicol environment. This phage-antibiotic combination is the only environment where phages remain potent even on day 2 of the experiment. We propose that the interaction between antibiotic and phage prevent the evolution of phage resistance. The mechanism behind this interaction remains to be uncovered.

### Mutants resistant to phage-only treatment are biased towards deep rough LPS

In the phage-only environment, mutants with a deep rough LPS emerge at a higher proportion than expected based on the assumption that mutations occur in proportion to gene length. We observed that 86 % of all mutants display the deep rough LPS phenotype (compared to the expected 58 %, based on gene length) (**Figure 5B**). It is unclear why deep rough mutants are overrepresented, but one reason could be that mutation rates are not uniform throughout the genome. It is possible that there are many loci with high mutation rates in genes that cause deep rough phenotypes (mutational bias) (Moxon et al., 1994; Lind et al., 2019; Bertels et al., 2021). Similarly, deep rough LPS candidate genes may contain a higher number of mutational targets (i.e., loci that produce a phage-resistant LPS phenotype) than do rough LPS candidate genes. Alternatively, mutations in deep rough LPS genes may be more likely to produce robust resistance – the ability of the mutation to reliably result in the establishment of a colony – in the presence of phages alone.

While deep rough LPS mutants are more abundant – at least in *E. coli –* they are more susceptible to certain antibiotics (Nikaido and Vaara, 1985; Wang et al., 2015). This characteristic could potentially be exploited by complementing phage therapy with antibiotics that specifically affect bacterial strains with a deep rough LPS structure.

### LPS modifications affect susceptibility to chloramphenicol and, to a lesser degree, gentamicin

Chloramphenicol and gentamicin are similar in their mechanism of action (Lin et al., 2018) and resistance mechanism (Burns et al., 1989; Recht and Puglisi, 2001; Kehrenberg et al., 2005; El’Garch et al., 2007). Thus, the reason that chloramphenicol affects deep rough mutants differently may hinge on its physicochemical properties. The two antibiotics differ in their type of inhibition, hydrophobicity, and molecular size. Specifically, chloramphenicol is a bacteriostatic, hydrophobic molecule of 323 Da, while gentamicin is a bactericidal, hydrophilic molecule of 477 Da. The key differences are most likely hydrophobicity and size, because small hydrophobic molecules can diffuse through the LPS barrier, while hydrophilic molecules generally require active transport for cell entry (Nikaido, 2003).

The increased susceptibility of deep rough LPS mutants to hydrophobic antibiotics is in line with previous studies. Deleting deep rough LPS genes (*waaC*, *waaF*, *waaG* or *waaP*) increases membrane hydrophobicity and permeability (Wang et al., 2015), can destabilise the membrane, and increases the entry of small hydrophobic molecules (Nikaido and Vaara, 1985; Chang et al., 2010; Pagnout et al., 2019). Testing more antibiotics could reveal whether type of inhibition, molecular size, or hydrophobicity has the strongest effect on susceptibility.

### Host range extension experiments could make ΦX174 a potent therapeutic agent

ΦX174 is very specific in its receptor recognition: it infects only about 0.8 % of 783 *E. coli* human and animal commensal strains (Michel et al., 2010), which currently makes ΦX174 unsuitable for therapeutic purposes. However, the host range of ΦX174 can be expanded through evolution experiments (Romeyer Dherbey et al., 2023; Romeyer Dherbey and Bertels, 2024) or genetic engineering (Yehl et al., 2019). Once the limitation of phage entry is overcome, ΦX174 can replicate its DNA and lyse hosts successfully (including various *E. coli* strains and other species such as *S. enterica* and *Enterobacter aerogenes*) (Suzuki et al., 1974; Mayr et al., 2005; Orta et al., 2023). If the host range of ΦX174 is extended, then the extensive knowledge of ΦX174 biology accumulated over the last few decades could be used in attempts to convert ΦX174 into an efficient therapeutic agent, especially when used in combination with an antibiotic to exploit shortened LPS structures (Romeyer Dherbey and Bertels, 2024).

More generally, our results show that phages used in combination with low levels of certain antibiotics can reduce the evolution of phage resistance in bacteria. This result indicates that the continued use of antibiotics may be beneficial even when the target bacteria are antibiotic-resistant. Of course, further studies are required to test whether our results also hold for higher, more clinically relevant antibiotic concentrations and resistance rates.

LPS modifications may also reduce pathogen virulence when treating with LPS-targeting phages. LPS can increase virulence (Burns and Hull, 1998; Raetz and Whitfield, 2002; Pier, 2007) and elicit immune responses (Alexander and Rietschel, 2001). If the evolution of phage-resistance leads to the truncation of LPS structures, then developing resistance to phage infection may render the pathogen less virulent. Future experiments with clinically relevant antibiotics and phages should be conducted to allow us to understand which LPS-targeting phage-antibiotic combinations are most beneficial in therapeutic settings.

## Methods

### Bacteria and phage used in this study

*E. coli* C wildtype and phage ΦX174 (GenBank accession number AF176034) were provided by H.A. Wichman (University of Idaho, USA). The *E. coli* C wildtype used in this study differs from the NCBI *E. coli* C strain (GenBank accession number CP020543.1) by 9 insertions and 2 nucleotide replacements (see Romeyer Dherbey et al., 2023). *E. coli* C phage-resistant strains R1 through R35 were generated in a previous study (Romeyer Dherbey et al., 2023); all other phage-resistant strains were generated in this study.

### Media and growth conditions

Bacteria and phages were grown in Lysogeny Broth (LB, Miller) supplemented with 5 mM CaCl_2_ and 10 mM MgCl_2_ to (henceforth referred to as ‘supplemented LB’). Supplemented LB agar (1.5 %) was used to plate phages and bacteria. Plates were incubated for ∼17 hours at 37°C. Where appropriate, chloramphenicol or gentamicin was added at the concentration indicated. For bacterial overnight cultures, 5 mL supplemented LB in a 13 mL culture tube (Sarstedt) was inoculated with a single colony from a supplemented LB agar plate, and incubated for ∼17 hours (37°C, 250 rpm). Fresh ΦX174 stocks were prepared weekly, using *E. coli* C wildtype cultures in exponential phase as described previously (Romeyer Dherbey et al., 2023).

### Determination of antibiotic susceptibility of bacterial strains

#### Determination of MIC

Preliminary measurements of the antibiotic susceptibility of *E. coli* C R1 through R35 were obtained using ETEST® strips (bioMérieux, France) on supplemented LB agar plates, in order to narrow down the range of concentrations to be tested for MIC determination (data not shown). Overnight cultures of *E. coli* C wildtype and phage-resistant strains were diluted 10-fold in Ringer’s solution. 50 µL of each diluted culture (∼5 x 10^6^ cells) was spread, using ∼15 glass beads (4 mm diameter), on a 90 mm supplemented LB agar plate (with or without antibiotic, as indicated). Plates were incubated at 37°C overnight (∼17 hours). The next day, plates were cooled to room temperature, and imaged for growth assessment. If bacterial growth was present on the images of all tested concentrations, a new round of testing with increased antibiotic concentrations was instigated. Tests were continued until there was no detectable growth across 3 replicates (CLSI, 2019).

#### Imaging to measure growth/susceptibility

Supplemented LB agar plates were uniformly photographed using a Chemidoc imaging system (Bio-Rad) under the colorimetric setting with an exposure of 1 second (see Data Availability statement). Images were analysed using OMERO (web server, version 5.14.1) to quantify growth (Allan et al., 2012). For uniform measurements, a rectangular region with a constant area (1157.85 mm², an area size free of light glare in most images) was marked on each image. Growth was measured in 8-bit pixel greyscale values from 0 to 255. Growth measures were obtained by subtracting the average greyscale value of a blank plate from the average greyscale value of each sample plate (**Supplementary Table S1**). A uniform bacterial lawn corresponded to greyscale values of 30 through 80 (depending on the thickness and density of the lawn). No, or very little, visible growth corresponded with greyscale values of 0 through 8. Average greyscale values of more than 8 are defined as “growth”, and values of less than 8 are defined as “no growth”. The sensitivity of the assay to the growth cut-off was tested by changing the growth cut-off to 12 in Model 3 (**Supplementary Table S7**). Median growth values across 3 to 5 biological replicates were used for **Figure 2**.

### Fluctuation test and isolation of phage-resistant *E. coli* C mutants

Phage-resistance rates were measured using a revised Luria-Delbrück fluctuation test (Luria and Delbrück, 1943; Rosche and Foster, 2000), across 9 plating environments (n=50 per environment): (set 1) ΦX174 alone; (sets 2-4) ΦX174 in combination with chloramphenicol (at 1 µg/mL, 2 µg/mL, or 6 µg/mL); (sets 5-7) ΦX174 in combination with gentamicin (at 2 µg/mL, 2 µg/mL, or 4 µg/mL); (set 8) 6 μg/mL chloramphenicol alone (exception, n=30); (set 9) 4 µg/mL gentamicin.

For each environment, 50 single *E. coli* C wildtype colonies picked from supplemented LB agar plates. Each colony was inoculated, in its entirety, into 100 μL of supplemented LB without antibiotic or phage (in 96-well plates). Inoculated cultures were incubated for 5 hours (37°C, 750 rpm). Cultures were resuspended by pipetting at the 3-hour and 4-hour timepoints to avoid settling from bacterial aggregation. After incubation, 50 μL of culture from each well was spread onto the corresponding supplemented LB agar plate. For all environments involving ΦX174, bacterial cultures were mixed with 50 μL of a high-titre phage lysate (∼3 x 10^8^ PFU/mL) prior to plating. Following initial incubation, plates with chloramphenicol at 6 μg/mL (with and without phage) were incubated further for 16 hours to check for slow-growing colonies.

We designed the experiment to be able to accurately measure rates from 10^-6^ per cell division to 10^-10^ per cell division. Specifically, the amount of bacterial culture to be plated on selection plates was altered to have a countable number of resistant colonies (Rosche and Foster, 2000; Zheng, 2017) (cite Zheng 2017, Rosche and Foster 2000). Using a pilot experiment, we determined the size of bacterial culture to be plated for the expected number of colonies to be above 1 but below 100. For environment (set 1), 50 μL of two-fold diluted bacterial culture was mixed with 50 μL of ΦX174 lysate; (sets 2-4) and (sets 5-7), 50 μL of undiluted bacterial culture (∼3 x 10^8^ CFU/mL) was mixed with 50 μL of ΦX174 lysate; for (set 8) and (set 9), 50 μL of 10-fold diluted bacterial culture was taken. The entire mixture was then plated on agar plates with the corresponding selective condition. For obtaining total CFU counts, cultures were serially diluted and plated on non-selective supplemented LB agar plates.

The next day, colonies were counted and one colony was randomly chosen from each replicate for which colonies were observed. Colonies were purified on non-selective supplemented LB agar plates, grown up in liquid culture, and stored at -80°C. In total, 222 mutants were stored in this way (ΦX174-only n=50; ΦX174+CL(1 µg/mL) n=41; ΦX174+CL(2 µg/mL) n=37; ΦX174+GM(1 µg/mL) n=46; ΦX174+GM(2 µg/mL) n=43; ΦX174+GM(4 µg/mL) n=1; ΦX174+CL(6 µg/mL) n=4).

### Resistance rate calculations and statistical analyses

Phage-resistance frequencies in *E. coli* C (i.e., the number of resistant colonies across 50 independent replicates) were used to calculate resistance rates. Resistance rates were calculated by the Lea-Coulson model following the method of Foster (Foster, 2006). This method was used because (i) the bacterial populations were each founded by a single cell, (ii) the model assumes that the growth rates of mutants and non-mutants are the same in the absence of selection, and (iii) the model allows for partial plating of cultures. The resistance rate and 95 % confidence interval were estimated using the Lea-Coulson (ε<1.0, i.e., partial plating) method of *webSalvador* (the web-application of R package *rSalvador*) (Zheng, 2017). Resistance rates were compared between different plating environments using the Lea-Coulson (ε<1.0) LRT from the R package *rSalvador* (RStudio version 2022.02.1) (RStudio Team, 2019; R Core Team, 2020).

### Whole genome re-sequencing

Samples were prepared for whole genome re-sequencing from 500 μl of overnight culture, using a DNeasy Blood & Tissue kit (Qiagen). Samples were sequenced at the Max Planck Institute for Evolutionary Biology (Plön, Germany) with a NextSeq 500, using a NextSeq 500/550 High Output Kit v2.5 (300 Cycles) kit (Illumina). Output quality was controlled using *FastQC* version 0.11.9 (Simon, 2010), read-trimmed using *Trimmomatic* (Bolger et al., 2014), and analyzed using *breseq* (Deatherage and Barrick, 2014). Aside from SNPs, other predicted mutations were confirmed by mapping raw reads to the corresponding modified genome in Geneious Prime (2023.1.1, https://www.geneious.com).

### Predicting phage-resistance rates

*Model 1*. The probability of a ΦX174-resistance mutation occurring in a specific *E. coli* C gene is assumed to be proportional to the length of the gene’s coding region (**Table 1**). We assumed that only genes previously observed (Romeyer Dherbey et al., 2023) confer resistance to ΦX174. In cases where a phage-resistant *E. coli* C mutant carried multiple mutations, only the mutation predicted to affect LPS structure was considered (e.g. *waaO* for R29, *galE* for R1).

The observed ability of phage-resistant mutants to grow in the presence of chloramphenicol or gentamicin was used to predict (i) phage-resistance rates, and (ii) genes in which mutations can confer resistance in phage-antibiotic combination environments (**Figure 2** and values in **Supplementary Table S1**). If, out of “n” mutants with a mutation in a gene, only “x” mutants grew in the presence of the antibiotic, (x/n) was assumed to be the probability that this gene confers resistance in a combination environment. Hence, phage-resistance caused by a mutation in such a gene is proportional to (x/n)* gene length (i.e., a weighted score). For example, out of five *galU* mutants, only one can grow at 2 µg/mL chloramphenicol. Thus, resistance caused by a mutation in *galU* (909 bp) has a weighted score of 182 (1/5*909) in ΦX174+chloramphenicol (2 µg/mL).

Resistance rate is proportional to the sum of all weighted scores of genes conferring resistance. The relative resistance rate for a combination environment is inferred as a ratio of the total weighted scores under each selective condition to that of the ΦX174-only environment (**Table 1**).

*Model 2*. The weighted scores assigned to each gene were changed to the number of mutations actually observed in the ΦX174-only environment (**Supplementary Table S6**). Mutations in two genes (*gmhB* and *galE*) are not observed in the ΦX174-only environment, meaning a 0 frequency, and 0 weighted score, in model 2. A weight of 0 incorrectly classifies these genes as not conferring phage-resistance, and will also predict 0 probabilities for combination environments. This issue was solved by adding a pseudo-count of 1 to all genes, including *galE* and *gmhB* (green highlight in **Supplementary Table S6**).

The inverse problem exists for *hldD* and *gmhA.* These genes were not observed in Romeyer Dherbey et al., hence the degree of antibiotic susceptibility incurred through mutations in these genes was not measured in **Figure 2**. *HldD* is a bifunctional gene and mutations can cause two distinct deep rough LPS phenotypes: *waaC*-like or *waaF*-like (Coleman, 1983; Schnaitman and Klena, 1993). Hence, phage-resistance rates and LPS type/gene distributions were calculated for both phenotypes. For final analyses, the model closest to the experimental observations (**Figure 2**) was used. *GmhA* mutants were assumed to grow similarly to *gmhB* mutant (i.e. growth in all environments except ΦX174+chloramphenicol (2 µg/mL)).

*Model 3*. The growth cut-off greyscale value was increased from 8-bit to 12-bit (see **Methods:** section “Determination of antibiotic susceptibility of bacterial mutants”). Candidate genes and weighted scores of the baseline were the same as in model 2 (**Supplementary Table S7**).

Note: When comparing the three models with each other, model 1 was modified to include *hldD* and *gmhA* to avoid the statistical problem of zero expectation for any category (see “model 1 (added genes)” in **Table 2**).

### Bootstrap analysis

A bootstrap analysis was performed to estimate sampling error in the 217 *E. coli* C strains isolated from phage-antibiotic combination environments (only 1 or 2 µg/mL antibiotic, hence 5 mutants from other environments were excluded). Mutants were re-sampled with replacement 100 times for each environment, in order to calculate the expected variation in the sampled observation. To re-sample genes in which mutations were not observed, but are nonetheless expected to confer phage-resistance when mutated, a pseudo-count of 1 was added to each gene that could provide phage-resistance in that environment (before sampling). The standard deviation of the 100 sampling events was used to draw error bars for both gene-wise and LPS-type distributions (**Figure 5** and **Supplementary Figure S5**).

### Co-evolution experiment

#### Environments

In a two-day co-culture experiment, we measured how cultures of *E. coli* C (and its resistant mutants) grow when exposed to phage-antibiotic combinations, compared to the presence of either phages or antibiotics alone. Growth was measured across ten environments: (sets 1-2) ΦX174 with chloramphenicol at 1 μg/mL or 2 μg/mL; (sets 3-4) ΦX174 with gentamicin at 1 μg/mL or 2 μg/mL; (set 5) ΦX174 only; (sets 6-7) chloramphenicol only, at 1 μg/mL or 2 μg/mL; (sets 8-9) gentamicin only, at 1 μg/mL or 2 μg/mL; (sets 10) no phage or antibiotic. We repeated the whole experiment twice and combined the data in the final plots, resulting in a total of 12 replicates for each environment (**Figure 6**).

#### Day 1

Six independent overnight cultures of *E. coli* C wildtype were used to inoculate six pre-cultures, by adding 100 µL of overnight culture to 5 mL supplemented LB. Once inoculated, the pre-cultures were incubated for 2 hours (37°C, 250 rpm) with loosened caps, to obtain growing cells. Next, co-evolution experiment cultures were set up in 96-well plates as follows. To avoid evaporation issues, the 36 wells around the edges of the plate were not used, and filled with 150 µL supplemented LB to serve as blanks/media controls. To the remaining 60 wells, in a randomized fashion (see **Data Availability**), a defined combination of bacteria, phage, and/or antibiotics were added (150 µL culture in total) see **Figure 6**) (n=6 per combination). To each well, 130 µL of either of the six bacterial pre-cultures was added. Phages were added at an approximate multiplicity of infection (MOI) of ∼0.25 to 0.3 (∼3*10^7^ CFU of *E. coli* C and ∼7.5*10^6^ to 9*10^6^ PFU of ΦX174), and antibiotics were added at concentrations of 1 µg/mL or 2 µg/mL (**Figure 6A**). The plate was incubated in a microplate reader (Agilent BioTek Epoch 2 Microplate Spectrophotometer) for 24 h (37°C, 365 rpm double orbital shaking). Growth was measured as absorbance at 600 nm (OD600) every 5 minutes.

#### Day 2

3 µL (2 %) of grown culture (from day 1) was transferred to 147 µL fresh supplemented LB in a new 96-well plate, with the same antibiotic conditions as before. Phages were not added separately (aside from the phages present in the 3 µL transfer). The plate was incubated under the same conditions for 24 h.

### Statistical testing

*T*-tests and Wilcoxon rank sum tests were used to determine whether there is a statistically significant difference in the mean concentration of antibiotic at which the growth of rough and deep rough *E. coli* C mutants is impeded (**Figure 2B**). *T*-tests were performed using the stats package of R (R version 4.2.0, RStudio version 2023.12.1) (RStudio Team, 2019; R Core Team, 2020). Wilcoxon rank sum tests, with an exact solution for tied datasets, were performed using the EDISON WMW calculator (Marx et al., 2016). Chi-squared tests for the goodness-of-fit of models (**Table 2**) were performed using the stats package of R (R version 4.2.0, RStudio version 2023.12.1) (RStudio Team, 2019; R Core Team, 2020). For the same, observed distributions of mutations by gene and by LPS type were compared with the distributions predicted by each model. In the two-day coevolution experiment, environments were compared using the Tukey’s Honest Significant Difference (HSD) test combined with ANOVA (**Figure 6B-C**). An ANOVA model was fitted to the area under the bacterial growth curves in each environment, followed by pairwise comparisons of the different environments using the Tukey’s HSD test, both using the stats package of R (R version 4.2.0, RStudio version 2023.12.1) (RStudio Team, 2019; R Core Team, 2020). Growth on the different days was compared separately.

### Data visualisation

Heatmaps (**Figure 2**) and histograms (**Figures 3B-C**, **4B**, **5**, **S2**, **S3**) were generated using Microsoft Excel (version 16.75.2). Schematic drawings (**Figures 1**, **2**, **3A**) were created using <u>BioRender.com</u>. The genomic distribution of mutation loci (**Figure 4A**) and boxplot of antibiotic susceptibilities (**Figure 2B**) were visualised using R (RStudio version 2022.02.1) (RStudio Team, 2019; R Core Team, 2020).

### Data availability

*Antibiotic susceptibility*. Agar plate images and raw greyscale values used to generate **Figure 2** and **Supplementary Table S1** are available in Edmond (doi: 10.17617/3.PIDVUT).

*Whole genome re-sequencing*. The *E. coli* C wildtype genome file (used as a reference), the ΦX174 wildtype genome (used as a reference), and raw sequencing reads of all sequenced phage-resistant *E. coli* C isolates are available in Edmond (doi: 10.17617/3.PIDVUT).

*Two-day coevolution experiment*. The 96-well plate-layout, growth curves for individual replicates and raw reads (growth data, source for **Figure 6**) for both days are available in Edmond (doi: 10.17617/3.PIDVUT).

## Acknowledgements

This work was generously supported by funds from the Max Planck Society (L.P.-F.B.). L.P. was supported by the International Max Planck Research School for Evolutionary Biology (IMPRS EvolBio). The authors also thank the Barrick Lab for their fluctuation test protocols, Prof. Qi Zheng for his help with rSalvador, Dr Carsten Fortmann-Grote for his help with image analysis, and all members of the Microbial Population Biology department who helped with experiments, especially Daniel Martens, Dr Elisa Brambilla, and Lena Zeller.

## Author Information

Affiliations: Max Planck Institute for Evolutionary Biology, Plön, Germany.

Contributions: L.P. and F.B conceived the study. L.P. and F.B. planned the experiments. L.P., N.R., J.R.D. and M.S. carried out the experiments. L.P. conducted the sequence analyses. L.P., J.G. and F.B interpreted the results. L.P. and F.B. wrote the original draft of manuscript, L.P., J.R.D., J.G., and F.B. reviewed and edited the manuscript. L.P. and J.G. prepared figures and tables. All authors reviewed and approved the manuscript.

## Ethics declaration

The authors declare no competing interests.

## Legends for Supplementary Material

**Supplementary Note S1.** Explanation of differences in antibiotic susceptibility profiles between isogenic bacterial strains in **Figure 2**.

**Supplementary Note S2.** Parallel evolution in the fluctuation test.

**Supplementary Note S3.** Predicted mutations from the fluctuation test that occur in genes not known to be related to LPS biosynthesis, assembly, or regulation.

**Supplementary Figure S1. Raw mean greyscale values measured for strains grown in the presence of chloramphenicol.** The figure shows mean greyscale values for growth of different *E. coli* C strains combined with different chloramphenicol concentrations. There are between one and three measurements for each combination shown on the x-axis. Each measurement is the mean greyscale value across a region of 1157.85 mm² in a photo of an LB plate inoculated with an *E. coli* C strain (see **Methods**). The black horizontal line at a greyscale value of 37 indicates the mode of the mean background greyscale value of blank LB plates without addition of bacteria. The green horizontal line shows the strict growth cut-off of 8 (37+8) and the red horizontal line shows the lenient growth cut-off of 12 (37+12). The colours of the datapoints indicate whether the strain is predicted to possess a rough or a deep rough LPS phenotype. Raw greyscale values and images are provided in Edmond (https://doi.org/10.17617/3.PIDVUT).

**Supplementary Figure S2. Raw mean greyscale values measured for strains grown in the presence of gentamicin.** For details see Supplementary Figure S1.

**Supplementary Figure S3. Mean greyscale values of strains grown in the presence of chloramphenicol (A) or gentamicin (B), clustered by whether the strain is rough or deep rough.** The figure shows the mean greyscale values for the growth of *E. coli* C at different antibiotic concentrations on LB plates for all strains shown in **Figure 2**. Each measurement is the mean greyscale value across a region of 1157.85 mm² in a photo of an LB plate inoculated with an *E. coli* C strain. The black horizontal line at a greyscale value of 37 indicates the mode of the mean background greyscale value, that is, of blank LB plates without addition of bacteria. The green horizontal line shows the strict growth cut-off of 8 (37+8) and the red horizontal line shows the lenient growth cut-off of 12 (37+12). The colours of the datapoints indicate whether the strain is predicted to possess a rough or a deep rough LPS phenotype. Raw greyscale values and images are provided in Edmond (https://doi.org/10.17617/3.PIDVUT).

**Supplementary Figure S4. Types of mutations identified across phage-resistant *E. coli* C strains isolated from the fluctuation test.** Bars show all predicted mutations grouped by mutational class (as % of total mutations “n” identified in each of the five plating environments sampled). If an isolate had two mutations, they were classified and counted separately. Insertions and deletions are further separated into four categories: (i) transposon insertion, where the inserted bases are a transposable element, (ii) duplication, where the inserted bases match the sequence preceding the mutated locus (up to 1.3 kbp), (iii) large deletion, where a region of more than 100 bp is deleted, and (iv) small indels, other insertions and deletions of <100 bp. Mutation details are provided in Supplementary Table S4. CL = chloramphenicol, GM = gentamicin.

**Supplementary Figure S5. Comparison of model 3 predictions with observed phage-resistance rates.** (**A**) Phage-resistance rates for phage-antibiotic combinations, relative to the phage-resistance rate in the ΦX174-only environment. Observed rates (black) are only qualitatively in line with the predicted rates from model 1 (gene length model, dark red; see **Table 1**). Rates predicted by model 2 (dark yellow; derived in **Supplementary Table S6**) are within 95 % confidence intervals for ΦX174+gentamicin environments, and are 2 % away from the 95 % confidence interval in ΦX174+chloramphenicol (2 µg/mL) environment. Phage-resistance rates for model 3 (blue, **Supplementary Table S7**) remain largely unchanged (relative to those for model 2), except for the ΦX174+gentamicin(2 µg/mL) environment (where the prediction becomes worse). (**B**) LPS phenotype distribution in phage-resistant *E. coli* C mutants isolated in the fluctuation test. Graphs show proportion of mutants with a certain LPS phenotype (values in **Supplementary Table S5**). Model 1 predictions (dark red) are qualitatively in line with predictions. In model 2 (dark yellow), LPS phenotype distribution in the ΦX174-only environment is now in line with the observations. Model 3 predictions (blue) improve slightly for ΦX174+chloramphenicol (2 µg/mL) and ΦX174+gentamicin (2 µg/mL) environments. (**C-F**) Gene-wise distribution compared to model 3 predictions (and model 2 in C-D). In ΦX174+chloramphenicol (1 µg/mL), mutations are predicted in *waaF, gmhB, gmhA* and *rfaH*, but none are observed. Similarly, in ΦX174+gentamicin (2 µg/mL), mutations in *gmhB*, *gmhA*, *hldE* are predicted but not observed. Compared to model 2, only predictions at ΦX174+chloramphenicol (2 µg/mL) (*waaP, galU*) and ΦX174+gentamicin (2 µg/mL) (*gmhB*, *gmhA*, *hldE*) change in model 3. Error bars show the standard deviation for each LPS phenotype (**B**) or each gene (**C-F**) from 100 bootstrapping iterations (resampling with replacement, see **Methods**).

**Supplementary Table S1. Growth of *E. coli* C strains in chloramphenicol and gentamicin.**

*No antibiotic: Growth on supplemented LB plates without antibiotic. ^#^Values indicate mean pixel density in bacterial lawns, after subtracting mean background density. Source data for **Figure 2**.

**Supplementary Table S2. Resistance rates observed in the fluctuation test.** **Nt is the total number of cells in the culture. ^#^ε is the fraction of cells plated. ^§^We counted phage-resistant colonies from 50 biological replicates for each plating environment, which were used to calculate phage-resistance rates and 95 % confidence intervals (with rSalvador; see **Methods**). ^$^The likelihood ratio test statistic from rSalvador was used to determine whether each calculated phage-resistance rate is significantly different to that of the ΦX174-only plating environment. *Mutation rates and significance intervals for GM 4 µg/mL were calculated without counting plates that contained too many colonies. ^†^Resistance rate relative to the resistance rate in the ΦX17-only plating environment, calculated for combination plating environments with chloramphenicol (CL) or gentamicin (GM). Raw data (colony counts) are provided in **Supplementary Table S3**.

**Supplementary Table S3. Colony counts from the fluctuation test.** **Phage-resistant colonies were counted from 50 biological replicates for each fluctuation test plating environment. The resulting colony counts were used to calculate phage-resistance rates and 95 % confidence intervals (see Methods). *“Burst”: some plates contained areas with large numbers of tiny colonies (i.e., uncountable). ^#^“invalid”: Plate discarded due to contamination.

**Supplementary Table S4. Details of mutations predicted in the phage-resistant *E. coli* C strains isolated from the fluctuation test.** ^$^DUP indicates a duplication event. ^#^“2,611,543, +A”: indicates an insertion of “A” after nucleotide 2,611,543. **“2,611,295 - 2,611,303, INS (2,783,815 - 2,784,945 INV)”: indicates positions 2,783,815 - 2,784,945 were inserted (inverted) at position 2,611,303. The inserted nucleotides are flanked by 2,611,295 - 2,611,303 on both sides (i.e., 2,611,295 - 2,611,303 are duplicated). ^§^E187fs_Ter#192 indicates a frameshift starting at E187, with a stop codon (Ter) generated at codon #192. F366mod_Ter#383 indicates protein sequence modified, e.g., due to a large insertion/duplication starting from F366, and (premature) stop codon (Ter) generated at codon #383. Difference between frameshift (fs) and modification (mod) is whether the original sequence of the gene is used as coding region with only a frameshift, or the new inserted sequence is used until the stop codon. ^||^Predicted LPS phenotypes for the mutants and the mutations in their genome. *Where multiple LPS genes carry mutations, the star indicates the gene predicted to cause the most dramatic LPS phenotype change. ^Column indicates whether the mutated gene is related to LPS biosynthesis, assembly, or regulation. †Even though the amino acid is unchanged, the codon affected is the start codon ATG, hence the initiation of translation is expected to be inhibited. See **Supplementary Figure S4** for a summary of mutational classes.

**Supplementary Table S5. Observed and predicted distribution of mutations in LPS genes.** ^$^Mutations in *rfaH* or promoter regions of the *waa* operon can affect any number of inner or outer core genes, hence are in a separate category (*rfaH*\*). Source data for Figure 5.

**Supplementary Table S6. Details of Model 2 used to predict phage-resistance rates.** See **Methods** for model 2 assumptions. **New genes. Highlighted in green: 2 genes in which 0 mutations were detected among isolates from the ΦX17-only plating environment, due to which a pseudo-count or pseudo-weight of 1 was added to all genes (including these 2 genes).

**Supplementary Table S7. Details of Model 3 used to predict phage-resistance rates.** See **Methods** for model 3 assumptions.

