## Supplementary Materials for "Chloramphenicol and gentamicin reduce resistance evolution to phage ΦX174 by suppressing a subset of *E. coli* C LPS mutants"

**Supplementary Information**

**Supplementary Note S1**

*Explanation of differences in antibiotic susceptibility profiles between isogenic bacterial strains in Figure 2.*

*E. coli* C wildtype-derived, phage-resistant strains R27 and R31 are genetically identical (Romeyer Dherbey et al., 2023), yet slightly different gentamicin growth profiles were measured for these mutants (**Figure 2A**). Specifically, R27 growth is fully inhibited at 2 µg/mL gentamicin, while R31 growth is slightly above the detection threshold. R31 growth is fully inhibited at the next gentamicin concentration measured (3 µg/mL). This discrepancy could be accounted for by small variations – e.g. in bacterial growth or media preparation – if 2 µg/mL were very close to the MIC of the R27/R31 genotype. If so, we would expect to observe growth slightly under or above the detection threshold in different (biological or technical) replicates. Detection of growth would then depend on the particular threshold. Indeed, when the growth detection threshold is increased to an average grayscale value of 12 (from 8), both R27 and R31 have an MIC of 2 µg/mL for gentamicin (see **Supplementary Table S1**).

**Supplementary Note S2**

### Parallel evolution in the fluctuation test

Some parallel evolution was observed in the fluctuation test; some independent isolates carry identical (loss-of-function) mutations, both within and between plating environments. For example, an 18 bp deletion (positions 74,589 - 74,606) occurs in seven mutants (1 from ΦX174-only, 1 from ΦX174+CL(1 µg/mL), 4 from ΦX174+GM(1 µg/mL), 1 from ΦX174+GM(2 µg/mL). In each of *hldD* and *waaT*, two mutations each occur in different replicates in the same environment. Notably, reductions in mutational target size do not appear to lead to higher degrees of parallelism; the most parallelism is seen in ΦX174+GM(1 µg/mL) (where there no mutational target size reduction is expected). Specifically, 1 *hldE* mutation is observed three times, 1 *hldD* mutation is observed twice, and 1 *waaG* mutation occurs four times. An alternative explanation for parallelism may be mutational bias (i.e., genomic loci that, due to specific features, are more susceptible to mutation) (Lind et al., 2019; Bertels et al., 2021).

**Supplementary Note S3**

### Predicted mutations from the fluctuation test that occur in genes not known to be related to LPS biosynthesis, assembly, or regulation

Several of the phage-resistant *E. coli* C mutants from the fluctuation test carry mutations in genes that are not known to be involved in the biosynthesis, assembly, or regulation of LPS. These include:

*dosP* (ΦX174-only environment): adjacent genes *dosP* and *dosC* collectively regulate the level of cyclic diguanylate (c-di-GMP) in the cell. C-di-GMP promotes the production of exopolysaccharides and other substances related to the extracellular matrix (Laventie and Jenal, 2020). DosP is involved in c-di-GMP degradation, thus a loss-of-function *dosP* mutation is expected to increase c-di-GMP levels and thus promote the production of extracellular polysaccharides.

*yfcV* (ΦX174+CL(1 µg/mL)): *yfcV* encodes a fimbrial protein, which is a known virulence factor in uropathogenic *E. coli* (Spurbeck et al., 2012). Since this fimbrial protein can increase adhesion of bacterial cells to eukaryotic epithelial cells *in-vitro*, it may also contribute to adhesion in other environments (Korea et al., 2010).

*glpG* (ΦX174+GM(1 µg/mL)): *glpG* encodes an intra-membrane protease (Keseler et al., 2021).

*yidC* (ΦX174+GM(2 µg/mL)): *yidC* is an essential gene that encodes a membrane protein insertase, which is part of the SecDFyajC-YidC holotranslocon membrane insertase complex (Nouwen and Driessen, 2002). Phages can use YidC for insertion of phage coat proteins through the membrane (Kol et al., 2008).

**Supplementary Figures**

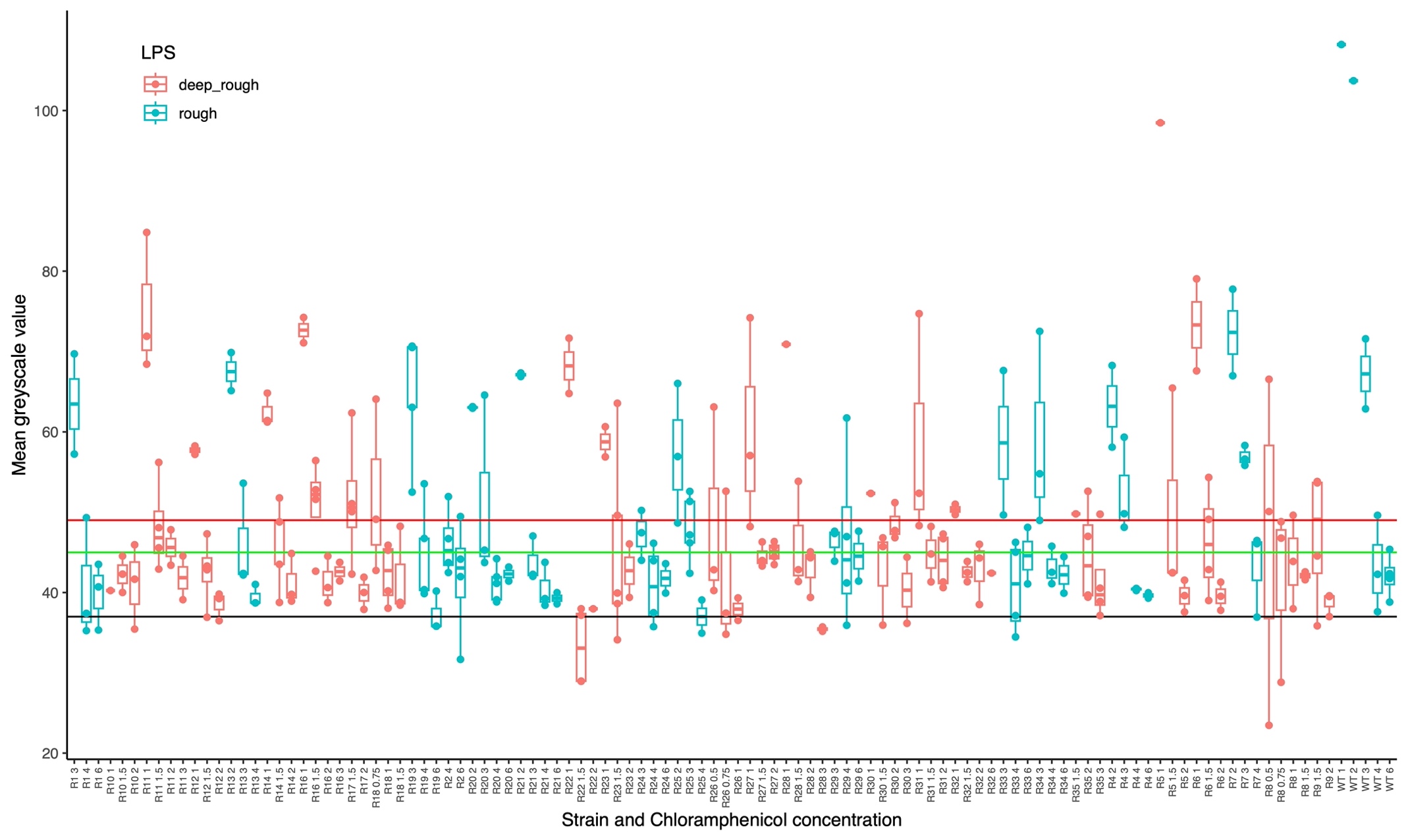

**Supplementary Figure S1. Raw mean greyscale values measured for strains grown in the presence of chloramphenicol.** The figure shows mean greyscale values for growth of different *E. coli* C strains combined with different chloramphenicol concentrations. There are between one and three measurements for each combination shown on the x-axis. Each measurement is the mean greyscale value across a region of 1157.85 mm² in a photo of an LB plate inoculated with an *E. coli* C strain (see **Methods**). The black horizontal line at a greyscale value of 37 indicates the mode of the mean background greyscale value of blank LB plates without addition of bacteria. The green horizontal line shows the strict growth cut-off of 8 (37+8) and the red horizontal line shows the lenient growth cut-off of 12 (37+12). The colours of the datapoints indicate whether the strain is predicted to possess a rough or a deep rough LPS phenotype. Raw greyscale values and images are provided in Edmond (https://doi.org/10.17617/3.PIDVUT).

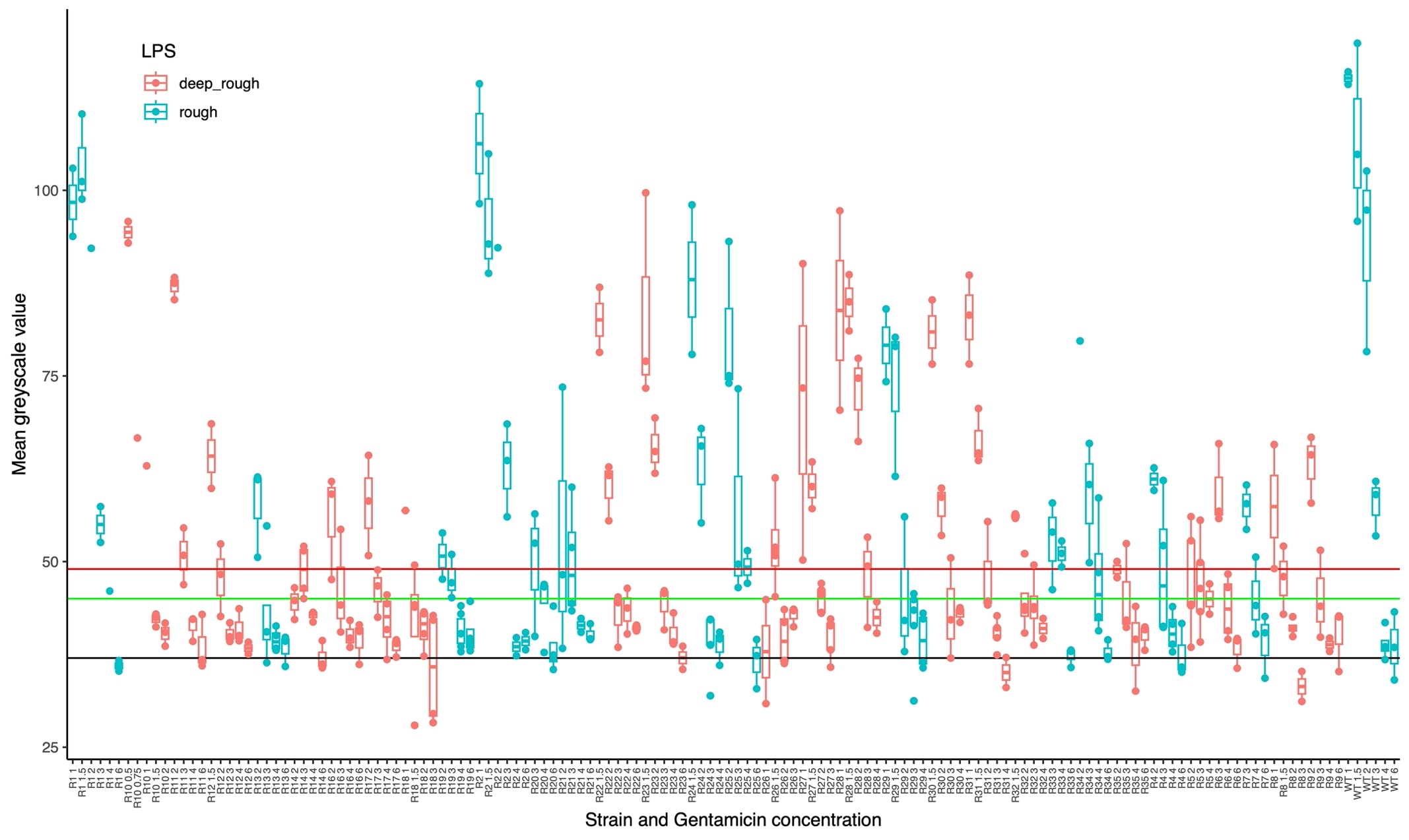

**Supplementary Figure S2. Raw mean greyscale values measured for strains grown in the presence of gentamicin.** For details see Supplementary Figure S1.

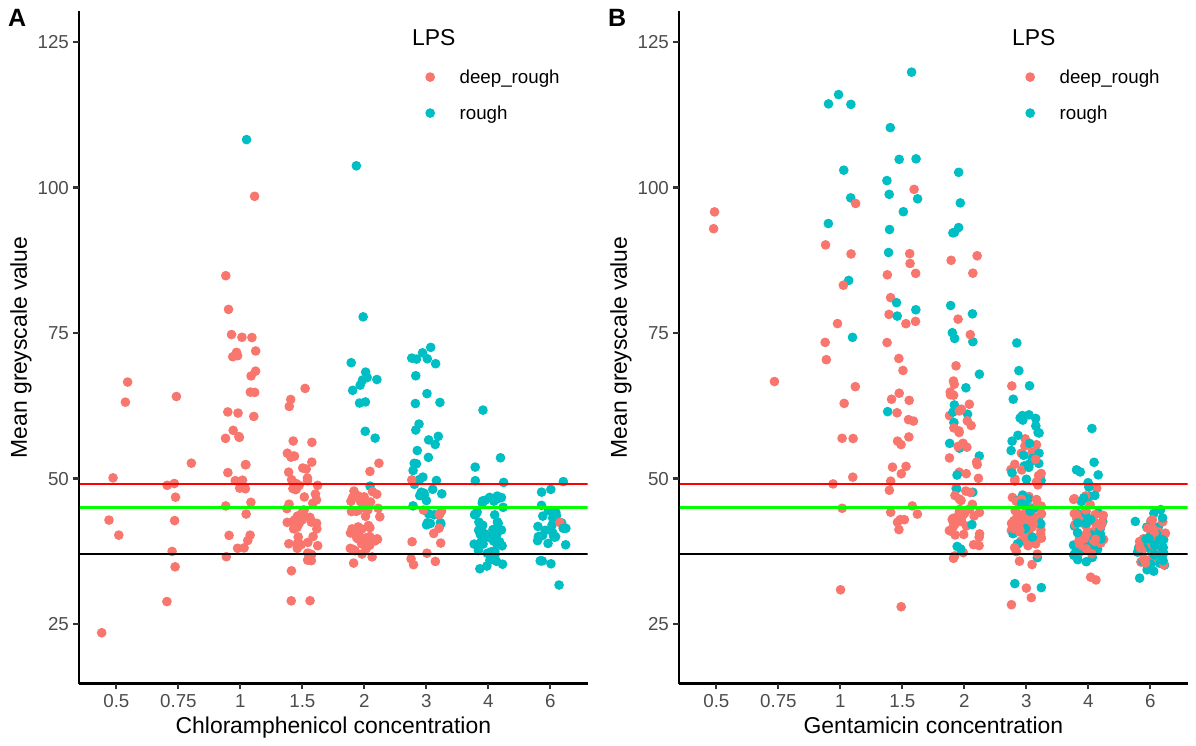

**Supplementary Figure S3. Mean greyscale values of strains grown in the presence of chloramphenicol (A) or gentamicin (B), clustered by whether the strain is rough or deep rough.** The figure shows the mean greyscale values for the growth of *E. coli* C at different antibiotic concentrations on LB plates for all strains shown in **Figure 2**. Each measurement is the mean greyscale value across a region of 1157.85 mm² in a photo of an LB plate inoculated with an *E. coli* C strain. The black horizontal line at a greyscale value of 37 indicates the mode of the mean background greyscale value, that is, of blank LB plates without addition of bacteria. The green horizontal line shows the strict growth cut-off of 8 (37+8) and the red horizontal line shows the lenient growth cut-off of 12 (37+12). The colours of the datapoints indicate whether the strain is predicted to possess a rough or a deep rough LPS phenotype. Raw greyscale values and images are provided in Edmond (https://doi.org/10.17617/3.PIDVUT).

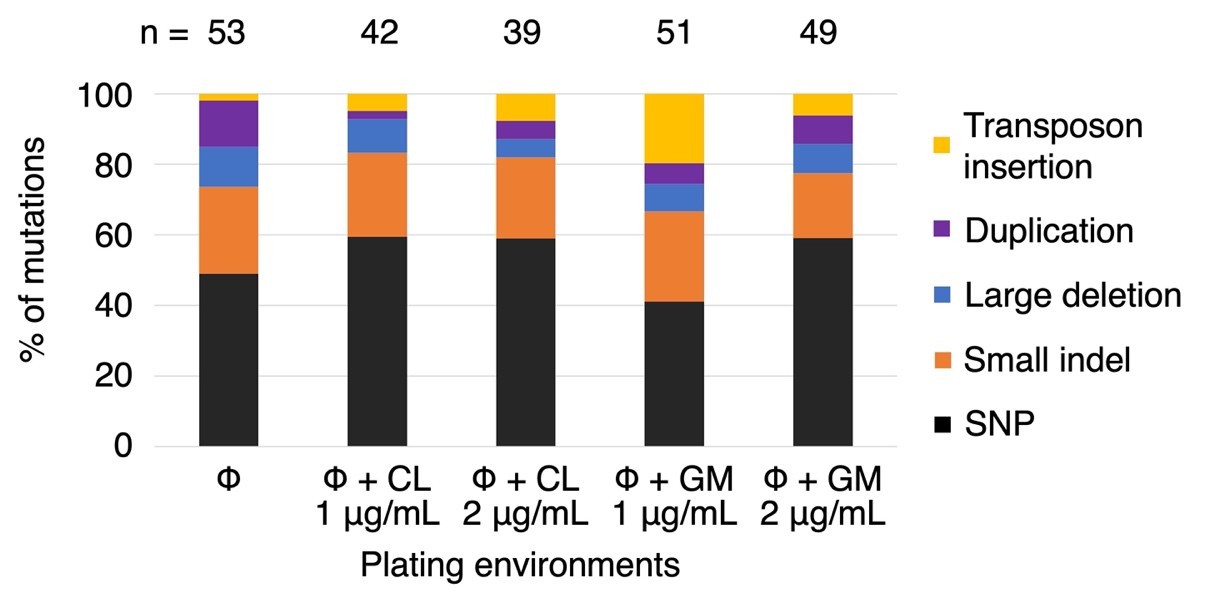

**Supplementary Figure S4. Types of mutations identified across phage-resistant *E. coli* C strains isolated from the fluctuation test.** Bars show all predicted mutations grouped by mutational class (as % of total mutations “n” identified in each of the five plating environments sampled). If an isolate had two mutations, they were classified and counted separately. Insertions and deletions are further separated into four categories: (i) transposon insertion, where the inserted bases are a transposable element, (ii) duplication, where the inserted bases match the sequence preceding the mutated locus (up to 1.3 kbp), (iii) large deletion, where a region of more than 100 bp is deleted, and (iv) small indels, other insertions and deletions of <100 bp. Mutation details are provided in **Supplementary Table S4**. CL = chloramphenicol, GM = gentamicin.

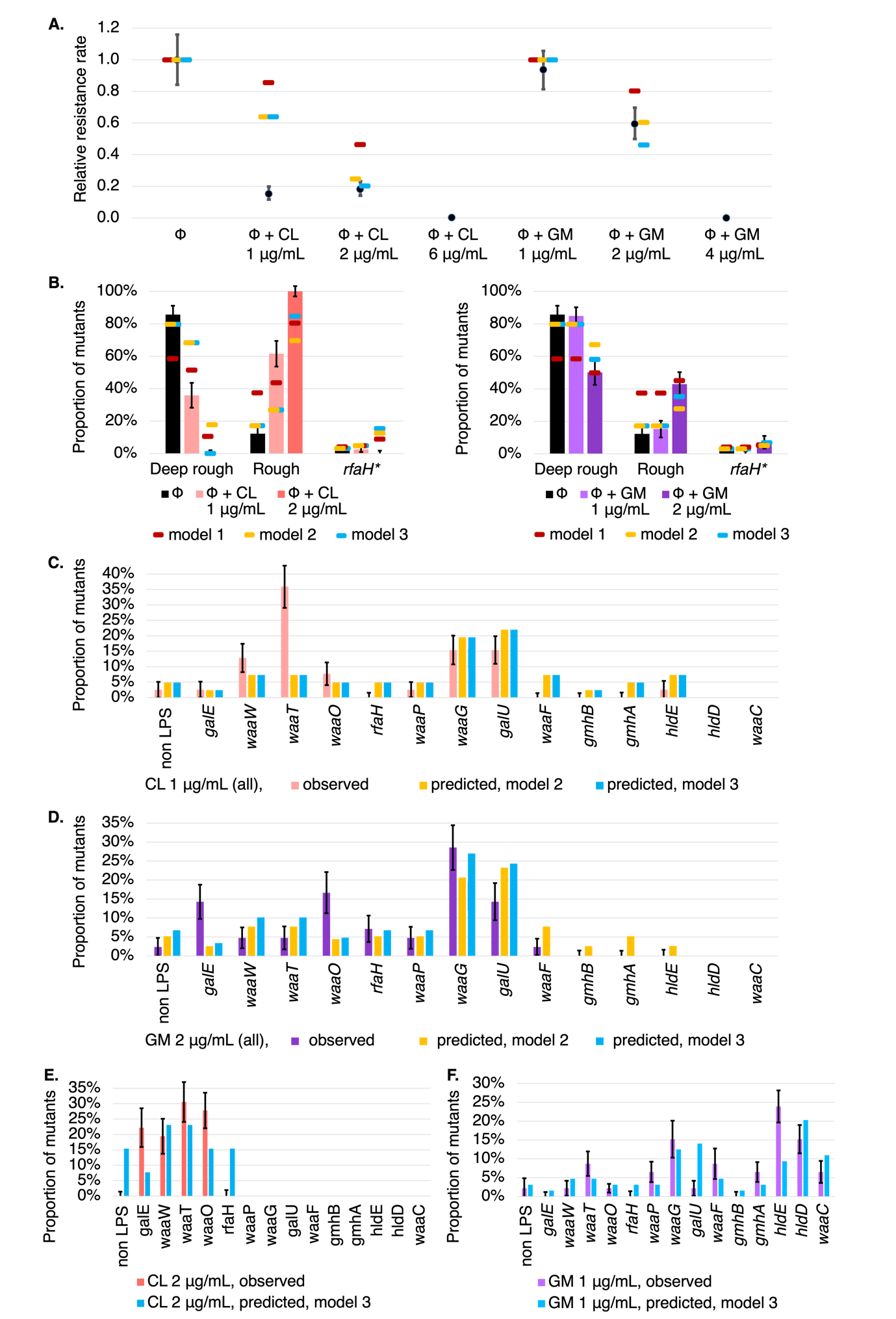

**Supplementary Figure S5. Comparison of model 3 predictions with observed phage-resistance rates.** (**A**) Phage-resistance rates for phage-antibiotic combinations, relative to the phage-resistance rate in the ΦX174-only environment. Observed rates (black) are only qualitatively in line with the predicted rates from model 1 (gene length model, dark red; see **Table 1**). Rates predicted by model 2 (dark yellow; derived in **Supplementary Table S6**) are within 95 % confidence intervals for ΦX174+gentamicin environments, and are 2 % away from the 95 % confidence interval in ΦX174+chloramphenicol (2 µg/mL) environment. Phage-resistance rates for model 3 (blue, **Supplementary Table S7**) remain largely unchanged (relative to those for model 2), except for the ΦX174+gentamicin(2 µg/mL) environment (where the prediction becomes worse). (**B**) LPS phenotype distribution in phage-resistant *E. coli* C mutants isolated in the fluctuation test. Graphs show proportion of mutants with a certain LPS phenotype (values in **Supplementary Table S5**). Model 1 predictions (dark red) are qualitatively in line with predictions. In model 2 (dark yellow), LPS phenotype distribution in the ΦX174-only environment is now in line with the observations. Model 3 predictions (blue) improve slightly for ΦX174+chloramphenicol (2 µg/mL) and ΦX174+gentamicin (2 µg/mL) environments. (**C-F**) Gene-wise distribution compared to model 3 predictions (and model 2 in C-D). In ΦX174+chloramphenicol (1 µg/mL), mutations are predicted in *waaF, gmhB, gmhA* and *rfaH*, but none are observed. Similarly, in ΦX174+gentamicin (2 µg/mL), mutations in *gmhB*, *gmhA*, *hldE* are predicted but not observed. Compared to model 2, only predictions at ΦX174+chloramphenicol (2 µg/mL) (*waaP, galU*) and ΦX174+gentamicin (2 µg/mL) (*gmhB*, *gmhA*, *hldE*) change in model 3. Error bars show the standard deviation for each LPS phenotype (**B**) or each gene (**C-F**) from 100 bootstrapping iterations (resampling with replacement, see **Methods**).
